# Learning Patterns of the Ageing Brain in MRI using Deep Convolutional Networks

**DOI:** 10.1101/2020.08.17.253732

**Authors:** Nicola K. Dinsdale, Emma Bluemke, Stephen M Smith, Zobair Arya, Diego Vidaurre, Mark Jenkinson, Ana I. L. Namburete

## Abstract

Both normal ageing and neurodegenerative diseases cause morphological changes to the brain. Age-related brain changes are subtle, nonlinear, and spatially and temporally heterogenous, both within a subject and across a population. Machine learning models are particularly suited to capture these patterns and can produce a model that is sensitive to changes of interest, despite the large variety in healthy brain appearance. In this paper, the power of convolutional neural networks (CNNs) and the rich UK Biobank dataset, the largest database currently available, are harnessed to address the problem of predicting brain age. We developed a 3D CNN architecture to predict chronological age, using a training dataset of 12, 802 T1-weighted MRI images and a further 6, 885 images for testing. The proposed method shows competitive performance on age prediction, but, most importantly, the CNN prediction errors Δ_*BrainAge*_ = *Age*_*Predicted*_ − *Age*_*True*_ correlated significantly with many clinical measurements from the UK Biobank in the female and male groups. In addition, having used images from only one imaging modality in this experiment, we examined the relationship between Δ_*BrainAge*_ and the image-derived phenotypes (IDPs) from all other imaging modalities in the UK Biobank, showing correlations consistent with known patterns of ageing. Furthermore, we show that the use of nonlinearly registered images to train CNNs can lead to the network being driven by artefacts of the registration process and missing subtle indicators of ageing, limiting the clinical relevance. Due to the longitudinal aspect of the UK Biobank study, in the future it will be possible to explore whether the Δ_*BrainAge*_ from models such as this network were predictive of any health outcomes.

**Highlights:**

- Brain age is estimated using a 3D CNN from 12,802 full T1-weighted images.
- Regions used to drive predictions are different for linearly and nonlinearly registered data.
- Linear registrations utilise a greater diversity of biologically meaningful areas.
- Correlations with IDPs and non-imaging variables are consistent with other publications.
- Excluding subjects with various health conditions had minimal impact on main correlations.

## 1. Introduction

The ageing of our global population is predicted to become one of the most significant social transformations of the twenty-first century [1]. This demographic shift has implications for all sectors of society, particularly healthcare. A range of neurodegenerative diseases, as well as cognitive decline, are challenges associated with this demographic change. In order to provide effective care and treatment, it is increasingly important to understand which individuals are most at risk of age-associated deterioration and how this deterioration will progress [2]. Even a healthy ageing process results in changes to the structure and function of the brain, and neuroimaging provides several methods of viewing these structural and chemical changes. The process of ageing is biologically complex and varied across the population, but neuroimaging modalities have been able to characterise general age-related changes in brain volume [3, 4, 5], cortical thickness [6] and white matter microstructure [7, 8]. Although there are general trends, the ageing process is both temporally and spatially heterogenous, and a large range of ‘healthy’ variation exists [3, 4, 5, 9, 10]. Distinguishing normal ageing from early stages of neurological disease is challenging, as the early stages of several disease states can resemble advanced ageing [3, 11, 12]. The ability to differentiate early signs of these diseases from normal ageing would hold great clinical value, as many current therapies are most effective with early intervention [13]. A better understanding of the healthy ageing process can help us to discriminate normal age-related changes from neurodegenerative diseases.

Several different approaches have been used to gain an understanding of the healthy ageing process. Most take the approach of developing a model of a healthily ageing brain and measuring how far a new patient differs from the model’s prediction of age: a greater divergence could indicate accelerated ageing, or possibly the onset of disease. It is first necessary to produce a model which accurately predicts the age of healthy subjects, in order to have a meaningful divergence measurement. To date, the most accurate methods have prediction errors in the range of 3.0-5.0 years, depending on the subject group [14]. There is little clinical relevance to predicting age from a medical image; if the subject can be scanned, it is probable we can ascertain their age by other means. However, error in age prediction in some models has been related to other physiological measurements of ageing, mortality, and general physical health [1, 15]. The fact that Δ_*BrainAge*_ = *Age*_*Predicted*_ − *Age*_*True*_ relates to cognitive performance, ageing fitness, and mortality [15, 16, 17] strongly supports the idea that Δ_*BrainAge*_ can provide clinically important insights [15].

The methods and type of neuroimaging data used to obtain the models in the literature vary widely; however, each uses Magnetic Resonance Imaging (MRI) data. There are multiple MRI modalities, each capturing different information about the tissue contents, structure, and function. The T1-weighted modality is the most commonly used because it is the most informative about the basic structure of the brain, especially the depiction of the main anatomical structures and tissues (i.e. grey matter (GM) and white matter (WM)) [18]. However, useful measurements can be obtained from other modalities: T2 FLAIR, susceptibility-weighted imaging (SWI), diffusion MRI (dMRI), or functional MRI (fMRI) images. Summary measures derived from the images (image-derived phenotypes (IDPs)) can be used for brain age modelling or, alternatively, voxelwise image data can be used. Although most of the prediction approaches have used IDPs derived from T1-weighted images [14, 19, 20, 21, 22, 23, 24, 25, 26, 27, 28, 29], combining IDPs from multiple image modalities (T1, T2 FLAIR, dMRI, fMRI) is also possible and has resulted in improved prediction performance [9, 21, 30]. Most reported methods extract IDPs and use them as input features for regression [25, 31], relevance vector machines (RVM) [19, 29, 32, 33, 34], support vector machines (SVM) [23, 26, 28, 32], or artificial neural networks (ANNs) [23, 24, 25, 31]. When the performance of ANNs have been compared to other types of machine learning such as SVMs [23], or Gaussian process regression [24], the ANN’s predictions yield the lowest mean absolute error (MAE) and highest correlation to the correct age. Although neural networks have previously been used in conjunction with IDPs, there has been a recent trend focussing on using convolutional neural networks (CNNs) to take advantage of data in both the extracted structural features and the raw, non-feature extracted images themselves [24], thus avoiding subjective decisions about which features are important. This not only reduces the amount of preprocessing, but also allows unknown relationships to be identified, as the model determines which features to use. This method has yielded amongst the lowest MAE scores reported, and, most notably, the Δ_*BrainAge*_ has correlated with patient mortality [15].

Age-related changes to the brain are subtle, nonlinear, and spatially and temporally heterogeneous, both within a subject and across a population [24]. Machine learning algorithms are particularly suited to address tasks with these properties and can produce a model that is sensitive to the large variety in healthy brain appearance. This can be more effective than comparing population ‘averages’ per age, which are unlikely to be representative of any single individual. Deep learning, specifically, has shown considerable promise for brain age prediction [24]. It can offer advanced ability to learn and represent features from the images, but deeper networks require more training data, which is often a limitation in medical imaging research. While machine learning methods have been used for this application for several years, the use of deep learning has been constrained due to sample size or computational ability. A large number of samples is needed to successfully train a deep neural network, and unlike the field of computer vision, ‘large’ neuroimaging databases often contain samples in the magnitudes of 10^2^s, not 10^6^s. In addition, although a CNN allows the whole image to be used, this is computationally very expensive.

Projects such as the UK Biobank and the Human Connectome Project are currently making unprecedented amounts of neuroimaging data accessible to researchers [35]. UK Biobank contains data from participants’ lifestyle, clinical measurements, genetic, and imaging data. The longitudinal aspect of the UK Biobank makes it well-suited for examining age prediction and health outcomes, as the Δ_*BrainAge*_ correspondence to later clinical outcomes will be measurable.

In this work, the power of CNNs and the UK Biobank dataset were employed to predict brain age. A 3D CNN architecture was developed, designed to predict age, and then tested for Δ_*BrainAge*_ correlation with other measurements from the UK Biobank. We used a training dataset of 12, 802 T1-weighted MRI images and a further 6, 885 images for testing. Although it is more computationally expensive, a 3D CNN preserves the structural information from all dimensions of the image, and has been shown to outperform 2D CNNs in some cases [36]. Since sex-specific effects have been demonstrated in both the ageing process and in previous prediction models [14, 37, 38], the 12, 802 subjects were separated by sex (6, 579 women and 6, 223 men) and considered separately at each stage of the experiments. This allowed examination as to whether the resulting Δ_*BrainAge*_ corresponded to different UK Biobank measurements for each sex. To investigate the performance of the network for subjects with a high age prediction delta, a subject with a high Δ_*BrainAge*_ was compared to an age-matched subject with a low Δ_*BrainAge*_. This provided an indication of whether the deltas held physiological relevance, or if they were simply evidence of the model’s failure to predict accurately. Since images from only one imaging modality in this experiment were used, the relationships between the Δ_*BrainAge*_ and IDPs from all of the imaging modalities in the UK Biobank were examined. The predictions were also compared to those derived from a second model that used only these multi-modality IDPs as training input, to give us insight into which modalities could be most useful for future age prediction models.

Through utilising a CNN we are able to create a model that has freedom to investigate spatial signatures, rather than relying on spatial priors or target structures, in contrast to other methods proposed to explore brain age in the UK Biobank. In both [39] and [40] image-derived phenotypes are used to train a linear model to predict brain age, with the former using the IDPs to create a single estimate of brain age and in the latter they explore whether multiple modalities can produce distinct modes that relate to different processes involved in ageing and disease. These methods allow us to explore different aspects of ageing using multimodal data but they require prior choice of the IDPs whereas our method is able to consider the whole of the input image and therefore there is potential for our method to pick up on different types of changes compared to the IDP-based models, especially those with subtle changes. These possible subtle changes are also a motivation for the exploration of the effect of registration on the model predictions. We do not fully understand all the nuances involved in the ageing brain, and therefore the exploration of different methods that consider different aspects of the problem using different approaches is important for further insight and understanding.

The main research goal of this work is, therefore, to explore the associations of brain age delta with the UK Biobank variables and by training with only the T1-weighted images we are able to explore the independent associations with the IDPs from the other modalities. Our goal is not to simply produce the lowest error on the age prediction task but to show independent biological associations with the deltas. This is therefore a contribution of this work beyond the current brain age literature on the UK Biobank. Furthermore, we explore the effect of the registration process on the predictions, and demonstrate that the most accurate models do not necessarily render the most biologically meaningful model and investigate what drives the models.

## 2. Methods

### 2.1. Data and Preprocessing

A total of 12, 802 T1-weighted structural MRI from the UK Biobank database [35] were used for training and a further 6, 885 for testing in this experiment, split by sex: the age distribution can be seen in Fig. 1. In line with existing findings about sex differences such as post-menopausal accelerated brain ageing [40], stratification by sex was also conducted in this study. The images were acquired at 1mm isotropic resolution using a 3D MPRAGE Sequence [18] and processed using the UK Biobank pipeline [41]. For these experiments, both the linearly registered and nonlinearly warped versions of the images (registered to the standard MNI template) were used to train the network separately, and the results compared. We trained on both linearly and nonlinearly registered images because, whilst part of many standard pipelines including the UK Biobank Pipeline [41], nonlinear registration to the MNI template suppresses morphological information. We know that these morphological changes are likely to be used by neural networks as a biomarker for age prediction and so it is important that we understand the effect of the registration process in the age predictions and Δ_*BrainAge*_ values calculated.

**Figure 1:**
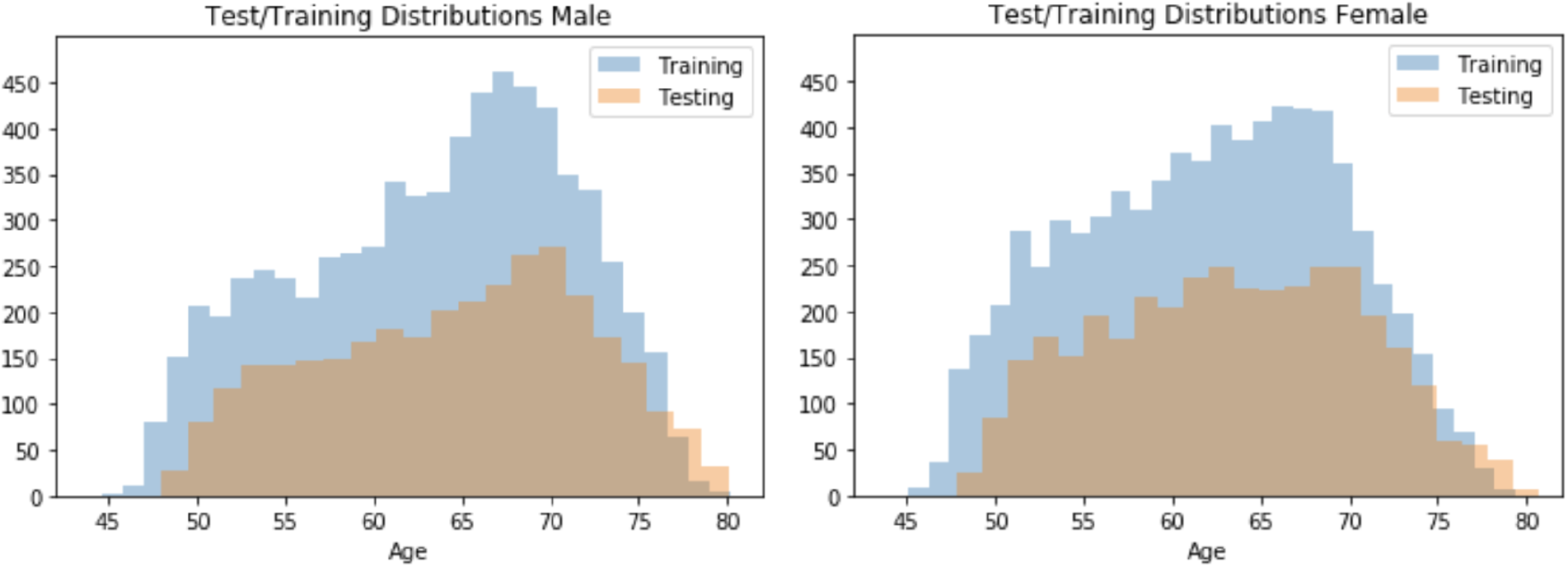
Training and test data histograms for male and female subjects. Maximum age = 80.6, minimum age = 44.6. Female mean (training) = 62.1, Female mean (testing) = 63.3, Male mean (training) = 63.4, Male mean (testing) = 64.6.

The original image dimensions were 182 × 218 × 182 voxels; however, to reduce the memory needed for computation, the *x* and *y* dimensions were resized to 128 × 128 voxels, and only 20 slices out of 182 were used, as can be seen in Fig. 2. To include a sufficient amount of anatomical information and resolution whilst reducing redundancy, every fourth slice was used from a region of 80 slices. The region was chosen so as to include as much of the brain as possible while still including some information on the cerebellum, whose structure and function is also impacted by ageing [42].

**Figure 2:**
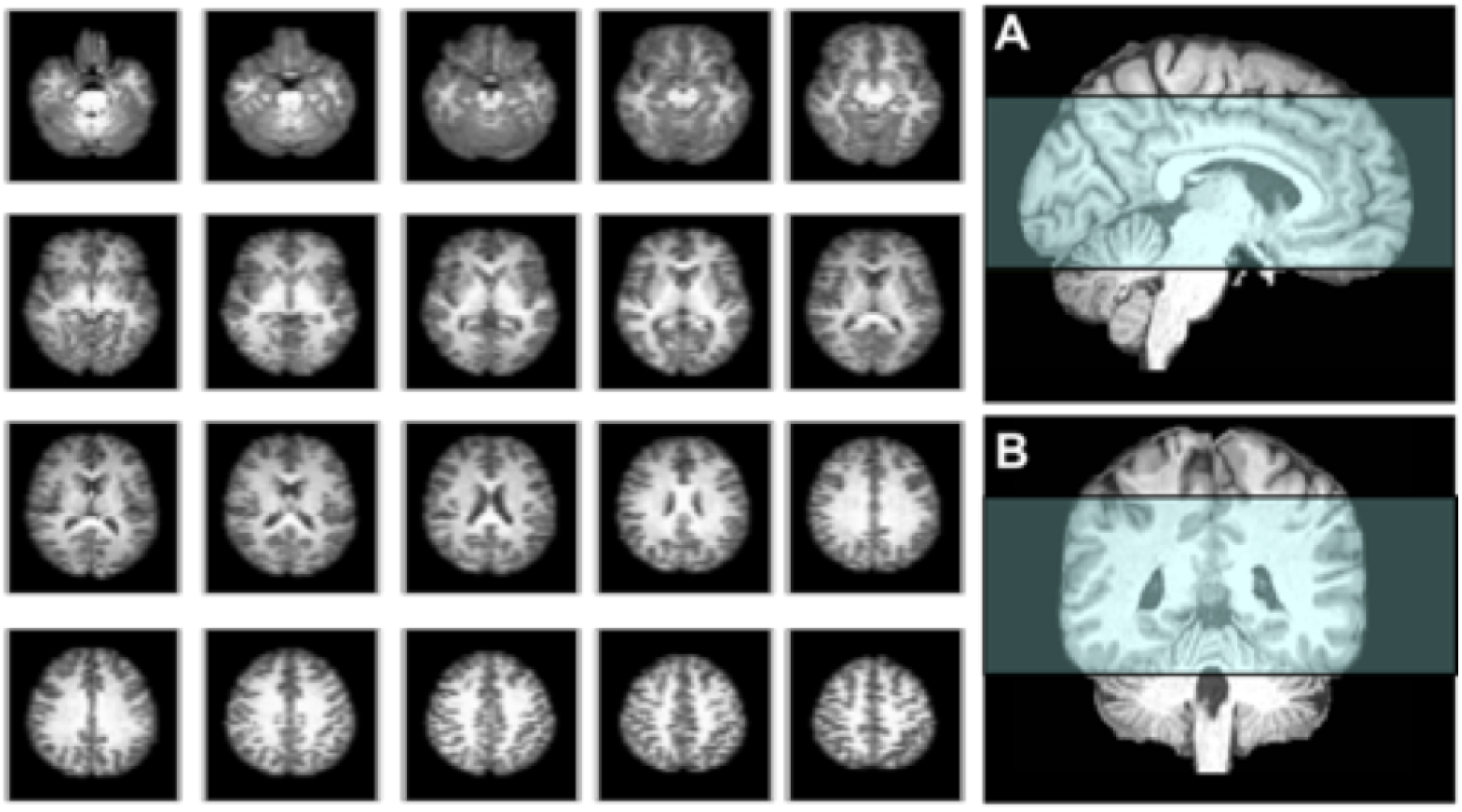
An example of structures contained in the axial view (128×128 pixels) of the 20 slices used. The region covered by the extracted slices is illustrated in the (A) sagittal and (B) coronal planes.

### 2.2. CNN Architecture

For this experiment, a deep 3D CNN was constructed. Sections of brain-extracted T1-weighted MRI scans were used as input to produce a single scalar regression output that represented the predicted age of the subject. Adapting the successful VGG-16 [43] architecture for age regression, the network used a mean squared error (MSE) loss function and consisted of 12 convolutional layers which convolved the input with a varied number of filters (*f*) of kernel size 3 × 3 × 3. The number of filters increased as the max-pooling decreased the spatial dimensions: this allowed the collection of a rich representation of features. The outputs of each convolutional layer were processed by a rectified linear unit (ReLU) activation function and then they were batch normalised [44]. The network contained four max-pooling layers with kernel sizes of 2 × 2 × 2 and a stride of 2. The output of the final convolution was then flattened and followed by two fully-connected layers of size 3*f* × 1 and *f* × 1, respectively. The final output layer of size 1 × 1 contained a linear activation function in order to predict age as the output scalar. To create the final network, three of these networks were combined to create an ensemble network, such that the final age prediction was the mean of the three network predictions. A complete schematic of both of these architectures is illustrated in Fig. 3.

**Figure 3:**
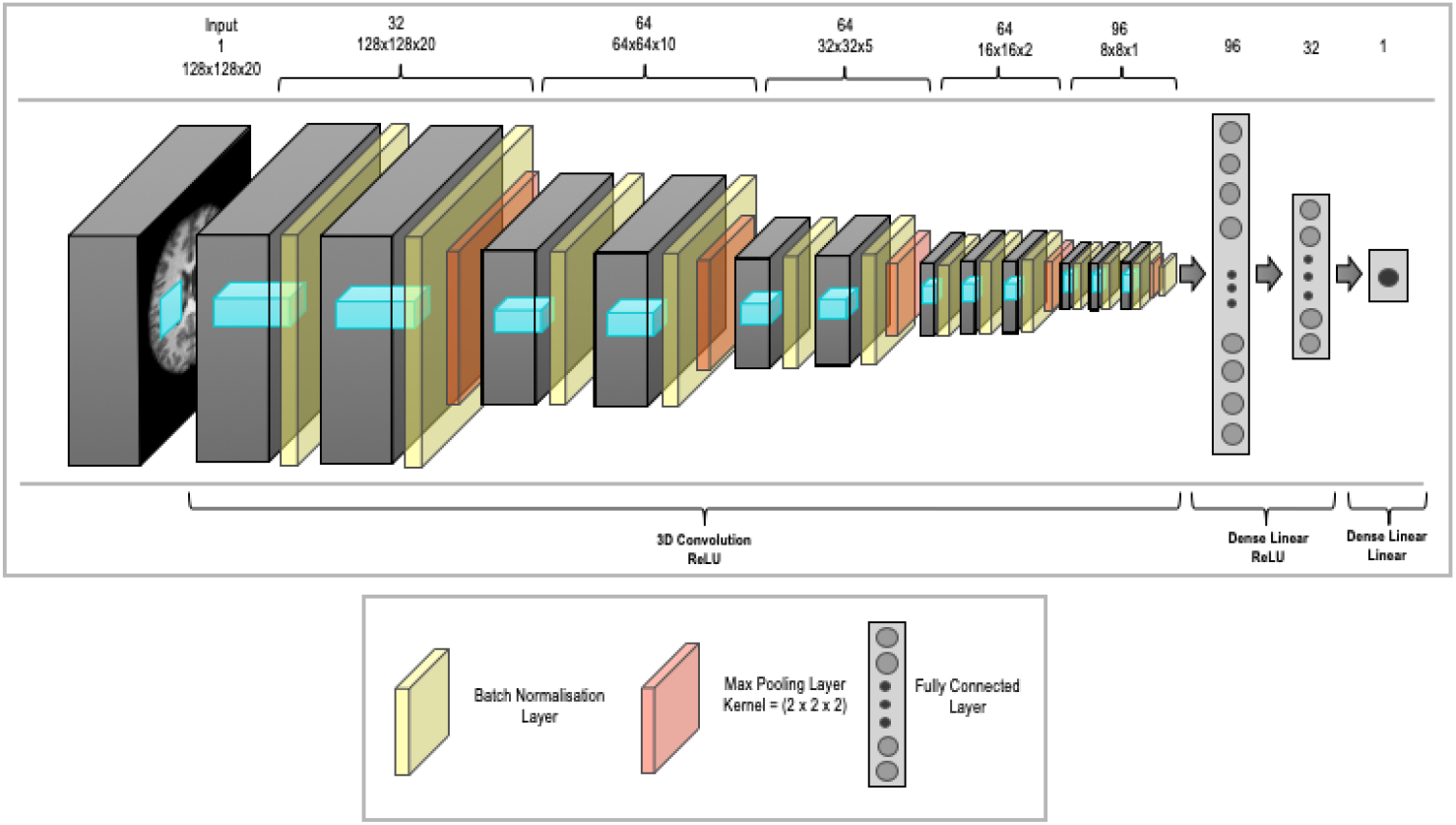
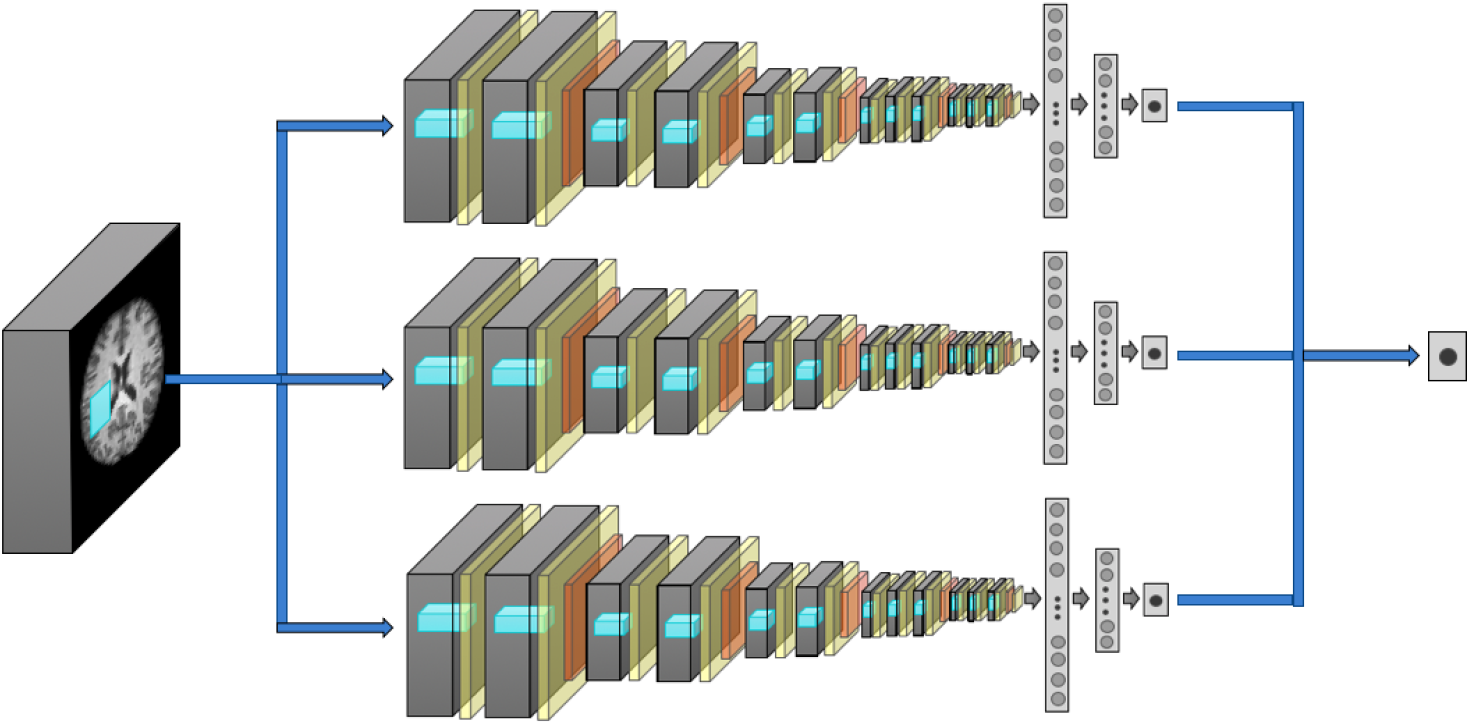
The network architectures used in this work. (a) The architecture of the fundamental network used in this work. As illustrated, each convolution is followed by batch normalization and ReLU activation. The number of filters in each of the twelve layers increases as f, f, 2f, 2f, 2f, 2f, 3f, 3f, 3f, 3f, 3f, 3f. (b) The ensemble network architecture. Each of the networks has the same structure but independent weights, learnt from randomly initialised weights. The aggregation is completed through averaging the outputs of the three individual networks.

### 2.3. Experimental Setup

A set of three networks were trained, for men and women separately, and the results were combined to form an ensemble network by finding the mean result. Ensembling was used because it increases the stability of the network predictions by reducing the variance. This, therefore, helps to make sure that the predicted Δ_*BrainAge*_ values are being driven by biological differences and not the random network initialisation or stochasticity in training.

The final ensemble networks were trained on the full dataset with a 90%/10% training/validation split and their performance was evaluated on a hold-out test dataset (subject numbers can be seen in Table 1). For each network and the final ensemble networks, the mean squared error (MSE), mean absolute error (MAE), and standard deviation of Δ_*BrainAge*_ = *Age*_*Predicted*_ − *Age*_*True*_ were calculated (the results can be seen in Table 2). Separate networks were trained and tested on the female and male subjects, as well as on the linearly registered and nonlinearly warped image datasets.

**Table 1:**
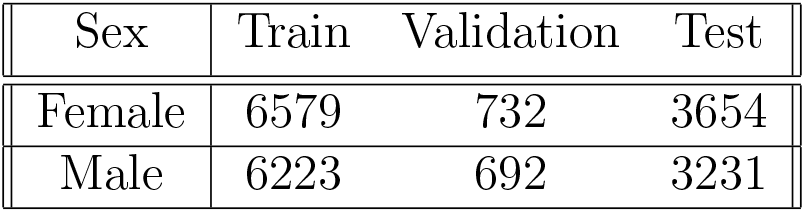
The number of subjects used in each step, separated according to sex.

| Sex | Train | Validation | Test |
| --- | --- | --- | --- |
| Female | 6579 | 732 | 3654 |
| Male | 6223 | 692 | 3231 |

**Table 2:** Ensemble Network results split by sex and image registration method.

|  | Linear Registered Input |  | Nonlinearly Registered Input |  |
| --- | --- | --- | --- | --- |
|  | Female | Male | Female | Male |
| MAE | 2.86±2.22 | 3.09±2.37 | 2.71±2.10 | 2.91±2.18 |
| MSE | 13.12±19.1 | 15.13±22.3 | 11.77±16.85 | 13.28±18.98 |
| R | 0.870 | 0.860 | 0.887 | 0.882 |

The training of the network was implemented using the Keras (v2.1.5) framework with a Tensorflow (v1.8.0) backend [45, 46] and performed on an NVIDIA Tesla P100 GPU. Optimization was achieved using the Keras RMSprop algorithm and mean squared error loss function over a maximum of 200 epochs. To minimize overfitting, the MSE on the validation set was monitored during training and the network was set to stop training if the validation MSE did not improve for 30 epochs. The initial learning rate was 10^−2^ and was set to decrease by a factor of 2 if the validation MSE did not decrease over 15 epochs, with a minimum learning rate of 10^−6^. In every epoch, the network was trained with a batch size of 16.

### 2.4. Correlation with Subject Biobank Measurements and IDPs

#### 2.4.1. Obtaining Δ_BrainAge_

The final predicted age was the output of the ensemble network, found by averaging the outputs of the individual networks in the ensemble. Once this mean age prediction was obtained, the difference between the network’s predicted age and true age was calculated with a simple subtraction: Δ_*BrainAge*_ = *Age*_*Predicted*_ − *Age*_*True*_. In this convention, a positive delta implies that the network predicted the subject’s brain to be older than the subject was, and a negative delta would imply the brain was predicted to be younger. Before performing any correlations, but after calculating MAE and similar performance metrics, consistent biases associated with linear and quadratic age were removed from the Δ_*BrainAge*_ acquired from the CNN.

#### 2.4.2. Relationship with Biobank Measurements

To examine the relationship between the CNN-derived deltas and UK Biobank measurements, univariate statistics were carried out using Pearson correlation analyses between the deltas and 8787 non-image derived variables from the UK Biobank database. These variables fell under various categories of lifestyle factors and physiological measurements and have been categorized in Miller et al. [18]. Sixteen of these categories were chosen to be of interest in our research, which include six lifestyle factors (Early Lifestyle Factors, Lifestyle General, Exercise, Work, Alcohol, and Tobacco), eight physiological or medical measurements (Physical General, Bone Density, Size, Cardiac, Blood Assays, Eye Test, Physical Activity Measures, Cognitive Phenotypes), and all variables contained in the Medical History and Mental Health Self-Report categories. The variables examined had all confounding variables removed (such as sex, age, head size, head motion, scanner table position, and imaging centre) and were then Gaussianised (quantile normalisation) [18]. The resulting p-values were sorted by category and plotted together with the Bonferroni correction and false discovery rate (FDR) correction thresholds (Fig. 7). The correlations were performed separately for each sex.

#### 2.4.3. Relationships with IDPs

To examine the relationship between the CNN-derived deltas calculated from T1-weighted images and the IDPs collected from all of the imaging modalities, univariate statistics were again performed using Pearson correlation analyses between the deltas and 941 IDPs as used by Miller et al. [18]. Confound variable removal, Gaussianization, and p-value calculation were all done in the same way as in Section 2.4.2.

#### 2.4.4. Comparison with IDP-Based Age Prediction

A separate model by Smith et al [39] for age prediction was used for comparison, where all of the IDPs, rather than images, were used to predict age. This was a linear regression model using all IDPs, a matrix of confounds, and a vector of ages as input variables. The model was regularised using an elastic net penalty, which is designed to promote sparsity in the model (i.e. to use a subset of the variables only), as well as leading the regression coefficients of correlated variables to have similar values. [47]. The elastic net dominates other regularisation choices, given an adequate choice of hyperparameters, which we selected using cross-validation. The prediction was then performed using multiple regression. Univariate pre-feature selection was completed [48] and only the 12.5% most correlated features were selected using nested five-fold cross correlation. In this paper, comparisons were made with all subjects for whom these features were available: 3212 female subjects were retained and 2717 male. Before performing any correlation between the IDP model-based deltas, the variance associated with age and quadratic age was removed following the method proposed by Smith et al (2019) [39]. A further explanation can be found in the supplementary material.

### 2.5. Ethics Statement

The UK Biobank has approval from the North West Multi-centre Research Ethics Committee (MREC) to obtain and disseminate data and samples from the participants, and these ethical regulations cover the work in this study. Written informed consent was obtained from all participants. Details can be found at www.ukbiobank.ac.uk/ethics. The UK Biobank was the only data used in this study.

## 3. Results

### 3.1. Age Prediction

The results of the CNN’s age prediction on both the female and male groups with both linearly registered and nonlinearly warped images can be found in Table 2. The resulting predicted ages have been plotted against the true ages for the hold-out testing data. For ease of comparison with other prediction methods in the literature, all MAE, MSE and SD for each network in the ensemble are reported in Table 3 and Table 4.

**Table 3:** Linear Registered Results for each network in the ensemble

|  | Female |  |  | Male |  |  |
| --- | --- | --- | --- | --- | --- | --- |
|  | Network 1 | Network 2 | Network 3 | Network 1 | Network 2 | Network 3 |
| MAE | 3.09±2.39 | 3.23±2.45 | 3.19±2.43 | 3.26±2.52 | 3.41±2.62 | 3.48±2.61 |
| MSE | 15.26±22.6 | 16.44±23.4 | 16.07±23.1 | 17.00±25.5 | 18.53±26.9 | 19.00±23.1 |
| R | 0.847 | 0.840 | 0.837 | 0.841 | 0.826 | 0.822 |

**Table 4:** Nonlinear Registered Results for each network in the ensemble

|  | Female |  |  | Male |  |  |
| --- | --- | --- | --- | --- | --- | --- |
|  | Network 1 | Network 2 | Network 3 | Network 1 | Network 2 | Network 3 |
| MAE | 3.09±2.37 | 3.17±2.42 | 2.87±2.22 | 3.32±2.56 | 3.23±2.41 | 3.21±2.47 |
| MSE | 15.15±22.0 | 15.88±22.8 | 13.15±19.0 | 17.06±27.8 | 16.27±23.0 | 16.42±24.2 |
| R | 0.847 | 0.844 | 0.869 | 0.839 | 0.849 | 0.844 |

When a high age prediction delta occurred for a subject, we compared their input images to a subject of the same age who had a low age prediction delta. In Figure 5, we demonstrate this comparison on Subjects 1 and 2: both subjects were 55 years old, but Subject 1 was predicted to be 72.31 with the linearly registered data and 68.28 with the nonlinearly registered data, while Subject 2 was predicted to be 55.44 with the linearly registered data and 58.43 with the nonlinearly registered data. In Figure 5B and D, the same slice linearly and nonlinearly registered, can be seen. Figures 5A and C show slices from the linearly registered brain for comparison between the two subjects.

**Figure 4:**
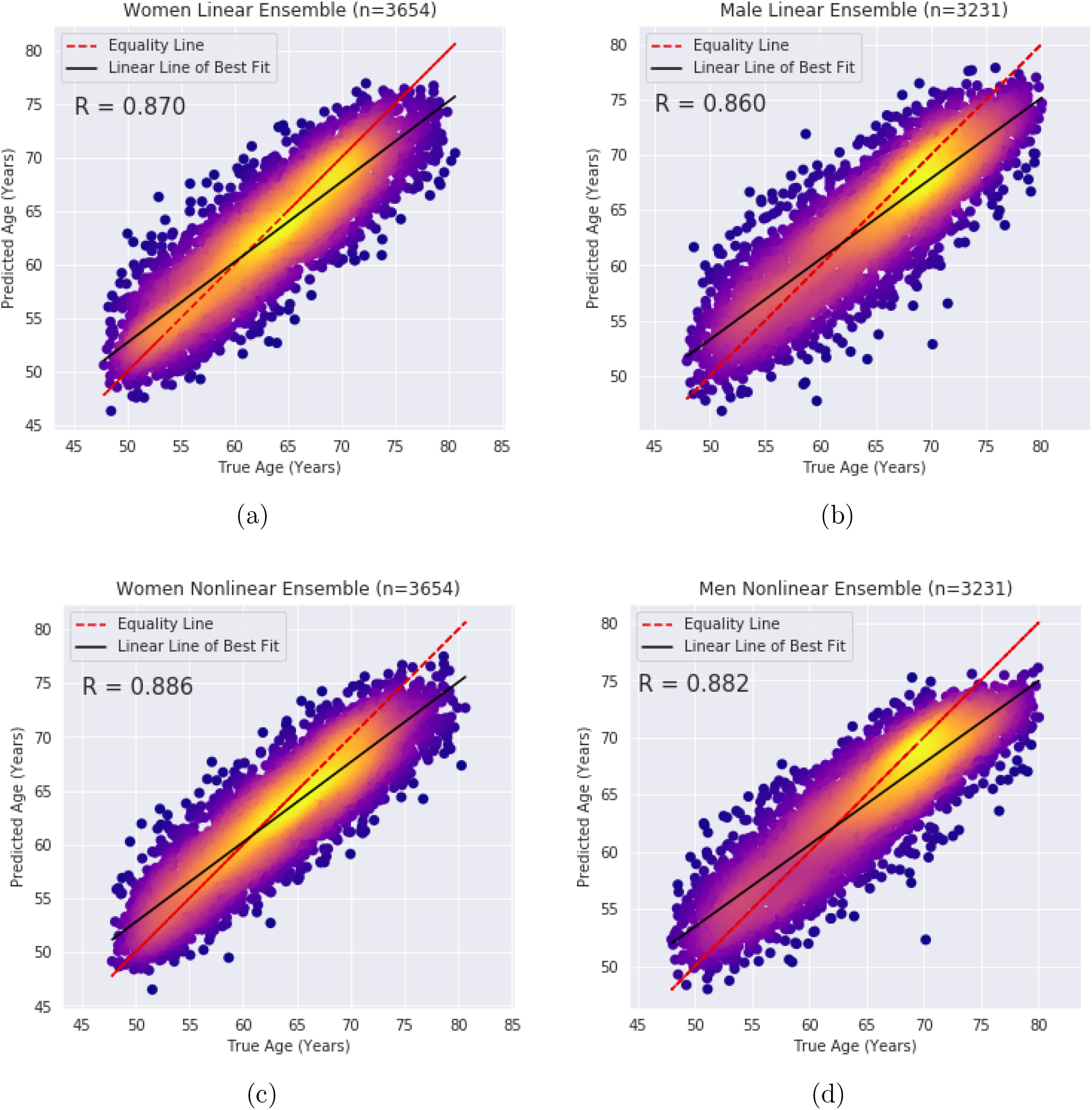
Density plot of the predicted ages vs the true subject age. a) predictions resulting from linearly registered images for the female subjects, b) predictions resulting from linearly registered images for the male subjects c) predictions resulting from nonlinearly registered images for the female subjects, d) predictions resulting from nonlinearly registered images for the male subjects.

**Figure 5:**
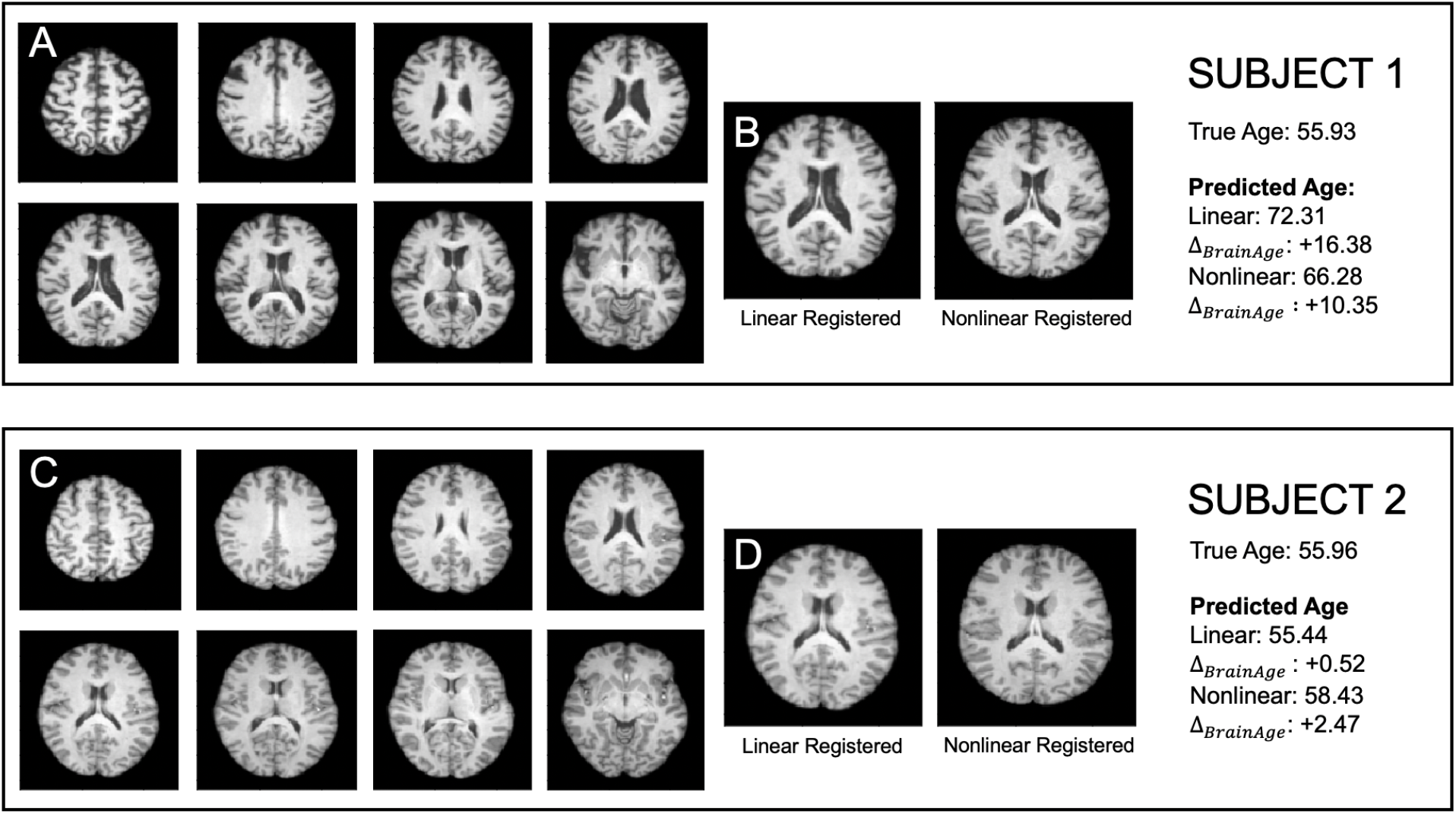
MRI Images from two subjects both with true age 55. A and C are example linear slices from the two subjects. B and D are the same slice for both subjects, both linearly and nonlinearly registered.

To examine which brain features are associated with subjects that have a higher predicted brain age, the non-linearly registered images of all subjects predicted to be 75-80 or 45-50 by the CNN in each sex group were averaged to form two mean composite images (Figure 6B). This process was repeated for the linearly registered images in Figure 6A. The largest difference was seen around the ventricles for both types of registration.

**Figure 6:**
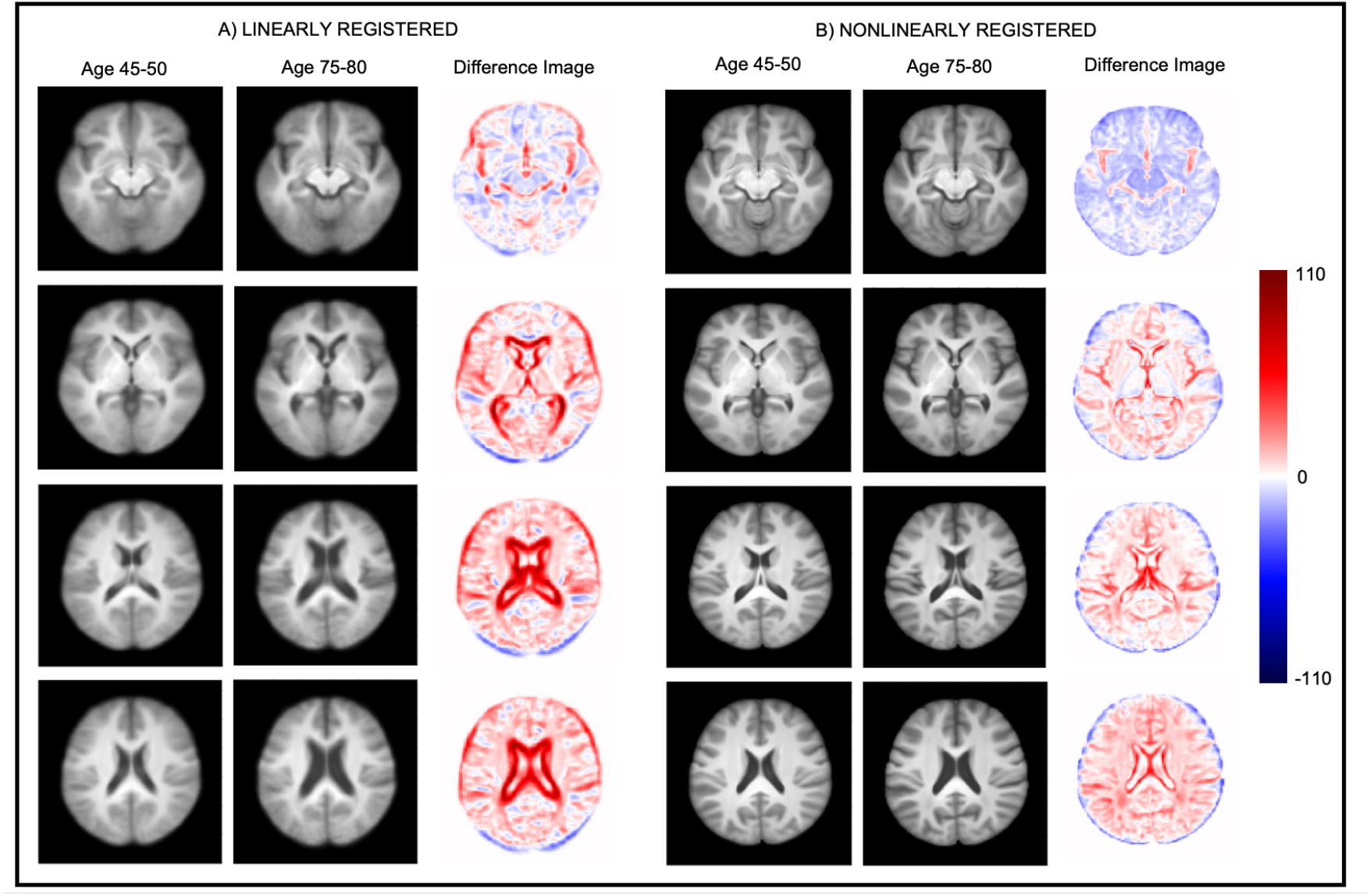
The mean of all the T1-weighted MRI of the subjects predicted by the CNN to be 45-50 and 75-80 in the female subject group: average of linearly registered images in (A) and nonlinearly registered images in (B). The male subjects showed similar changes. The four slices shown are sampled from the inferior (top row) to superior (bottom row) regions of the brain.

### 3.2. Relationships with UK Biobank Measurements

#### 3.2.1. UK Biobank Variables

The relationship between the deltas (obtained from the linearly warped images) and 8787 non-image derived variables from the UK Biobank are plotted in Figure 7. For the female group, using deltas from 3654 subjects, 14 variables passed the Bonferroni threshold including 4 variables included from the Medical History, 2 from Physical Measurements (Cardiac), 4 from Physical Measurements (Bone Density) and 4 from Physical Measurements (General). Similarly, for the male group, using deltas from 3231 subjects, 33 passed the Bonferroni threshold including 28 variables from the Medical History, 1 from Early Life Style (Food and Drink) and 4 from Physical Measurements (General).

**Figure 7:**
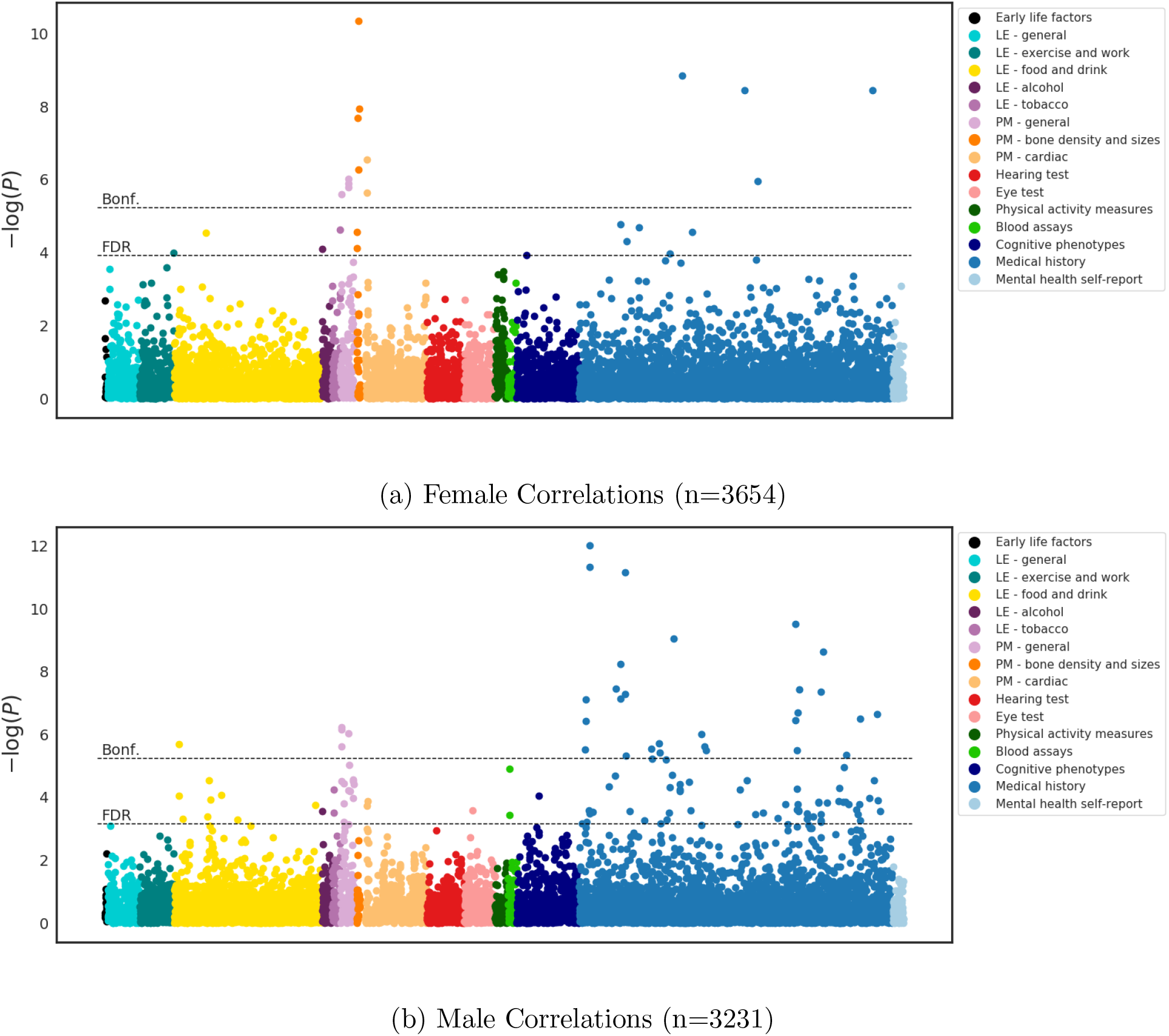
Manhattan plot relating each of the non-imaging UK Biobank variables with the delta measurements from the CNN (with linearly registered images as input) on female and male subjects. Each dot represents the statistical significance of the correlation between one UK Biobank variable and the deltas. A total of 8787 variables were tested, of which 14 and 33 passed the Bonferroni threshold for the female and male groups, respectively.

#### 3.2.2. Image Derived Phenotypes

Using the linearly registered images as input, 596 and 622 correlations passed the Bonferroni threshold in the female and male groups respectively. It must, however, be noted that the threshold for the T1-related IDPs is not strictly valid as the variables and CNN predictions were derived from the same source and thus are not independent. The relationship between the Δ_*BrainAge*_ and the 941 IDPs can be seen in Figure 8. To examine these more closely, the anatomical location of each Tract-Based Spatial Statistic (TBSS) IDP is illustrated in Figure 9. These TBSS IDPs are derived from white matter tracts and quantities calculated from diffusion MRI: fractional anisotropy (FA; the directional variation of water molecule diffusion), mean diffusivity (MD; the average magnitude of water molecule diffusion), intra-cellular volume fraction (ICVF; showing neurite density), isotropic volume fraction (ISOVF; showing extracellular water diffusion) and tract complexity/fanning (OD)[49].

**Figure 8:**
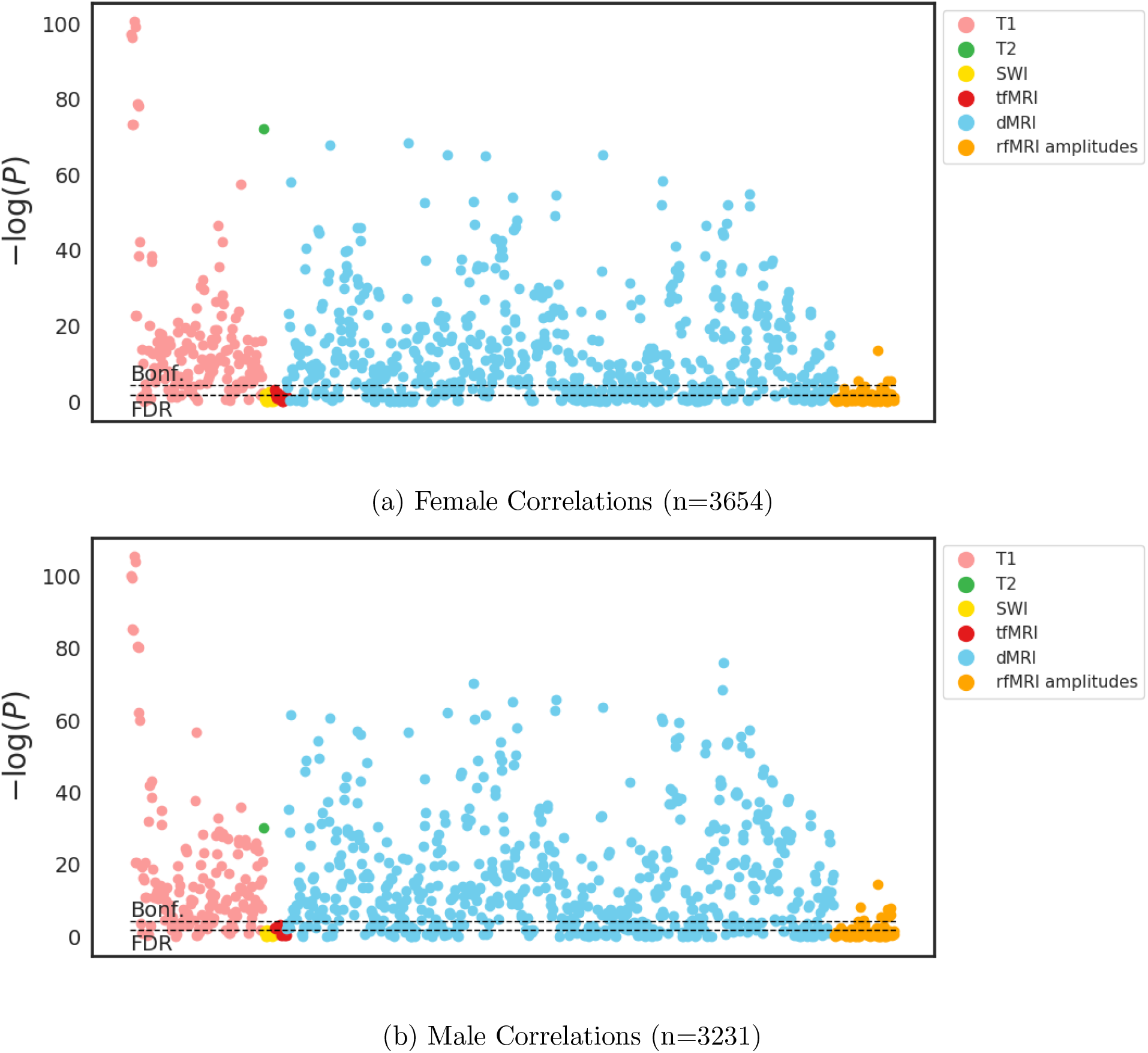
Manhattan plot relating each of the IDPs with the delta measurements from the CNN (with linearly registered images as input) on female and male subjects. Each dot represents the statistical significance of the correlation between one IDP type and the deltas. A total of 941 were tested, of which 596 and 622 passed the Bonferroni threshold for the female and male groups, respectively.

**Figure 9:**
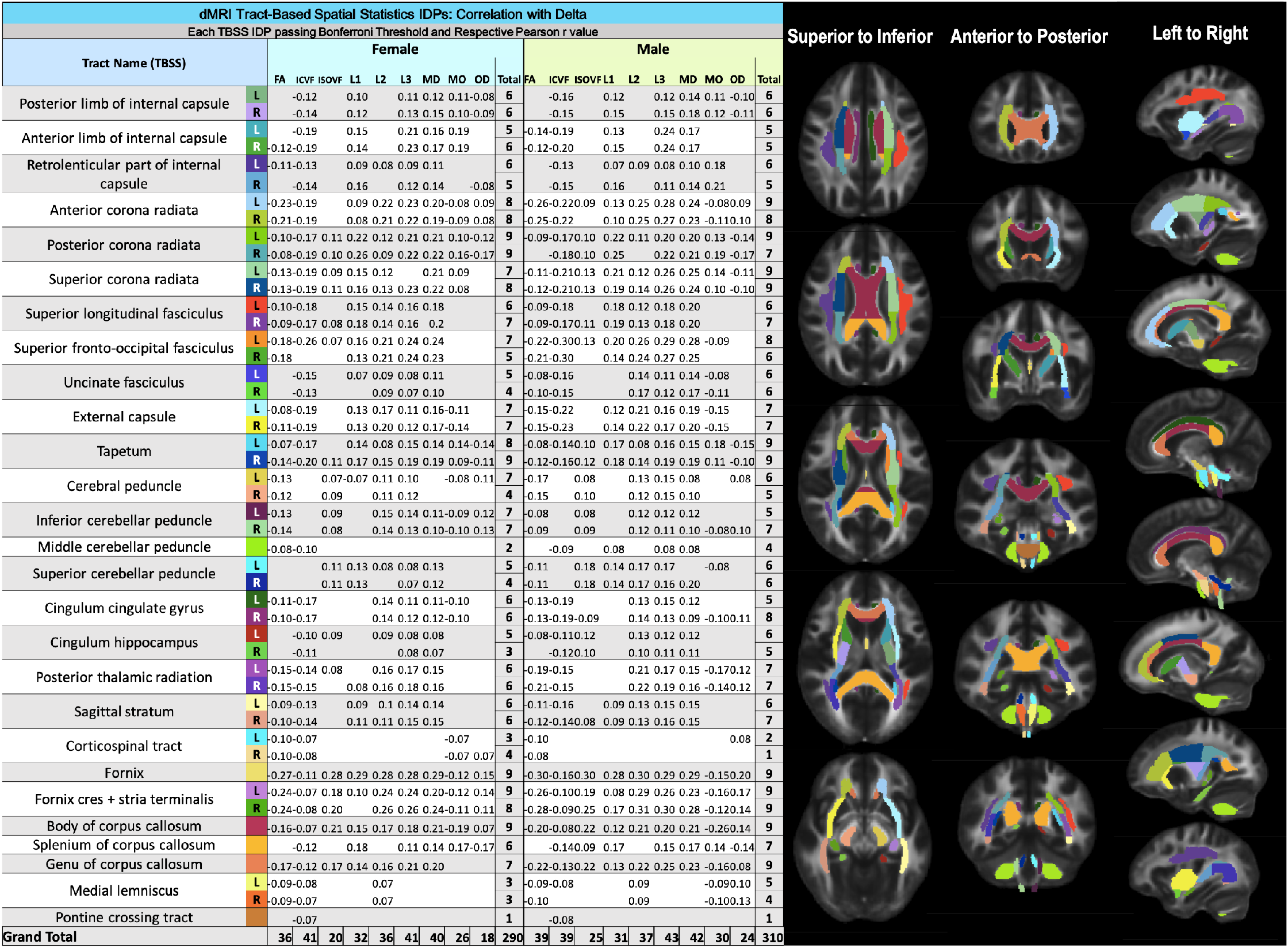
A chart of each IDP passing the Bonferroni threshold in Figure 8 and the corresponding correlation coefficients. The white matter tract associated with each listed IDP is displayed, based on the JHU white-matter tractography atlas [49] on top of the FMRIB58-FA standard-space FA template [50]. Acronyms: fractional anisotropy (FA), mean diffusivity (MD) intra-cellular volume fraction (ICVF); isotropic volume fraction (ISOVF), tract complexity/fanning (OD); L1, L2, and L3 are the three eigenvalues of the diffusion tensor.

To confirm the utilty of the network we also trained the network only on healthy subjects by two definitions: the first, the removal of conditions known to impact on the brain, and the second, the removal of these subjects plus those with conditions we have found to correlate with Δ_*BrainAge*_ – diabetes and hypertension – that had not already been removed. These were also found to correlate with Δ_*BrainAge*_ in [39]. Further details of conditions removed and the results can be seen in the supplementary material. The results were consistent across the experiments, showing that the results are not driven by subjects with known conditions.

### 3.3. Relationships with IDP-based Age Prediction

Illustrated in Figure 10, the CNN-derived age predictions (using linearly registered images) and IDP-derived age predictions were strongly correlated (female *r*=0.775, male *r*=0.791). The deltas from each model were also moderately correlated after being deconfounded for age (female *r*=0.409, male *r*=0.431). The CNN-derived age predictions using nonlinearly registered images (not shown) resulted in very similar correlations (female *r*=0.740, male *r*=0.799; deconfounded deltas: female *r*=0.399, male *r*=0.446). Although high correlations were found, the results from the two methods were not the same, as discussed below. The correlations with the IDPs and non-imaging variables can be seen in Figures 11 and 12. The Elastic Net achieved an MAE of 3.87 ± 2.88 on the female group and 4.15±2.99 on the male group compared to the CNN achieving 2.89±2.17 on the female group and 2.71 ± 2.09 on the male group on the same subset of subjects.

**Figure 10:**
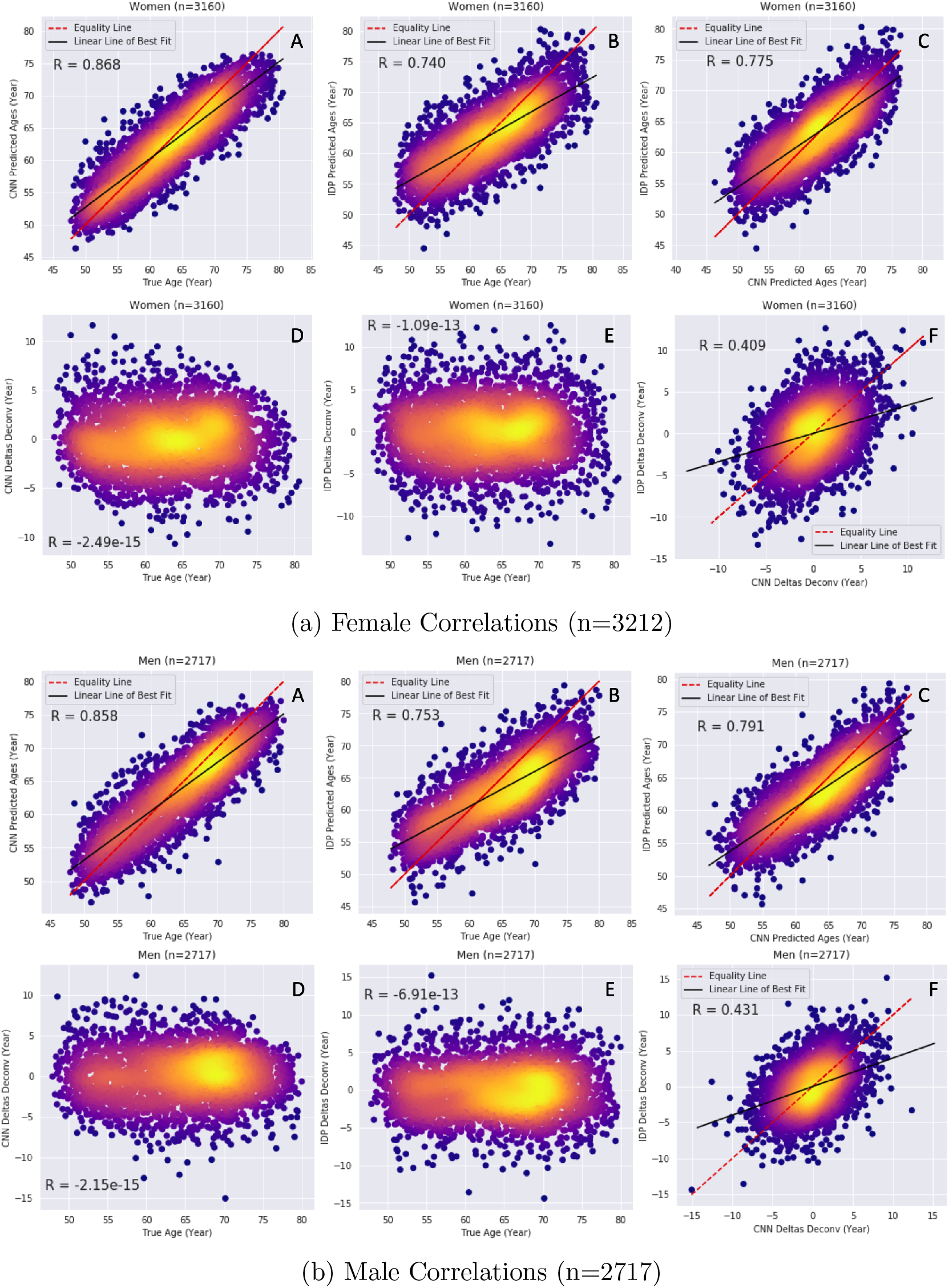
Density plots of the predicted ages from the CNN model (A) and IDP-based model (B) versus true age. (C) shows the relationship between the predicted ages of the CNN model and the IDP-based model. The deltas (deconfounded for age) from the CNN model and IDP-based model are plotted against true age in (D) and (E), respectively. (F) shows the relationship between the deltas from each model. The results are shown separately for each sex group. These results are based on the results from the CNN, trained on linearly registered images.

**Figure 11:**
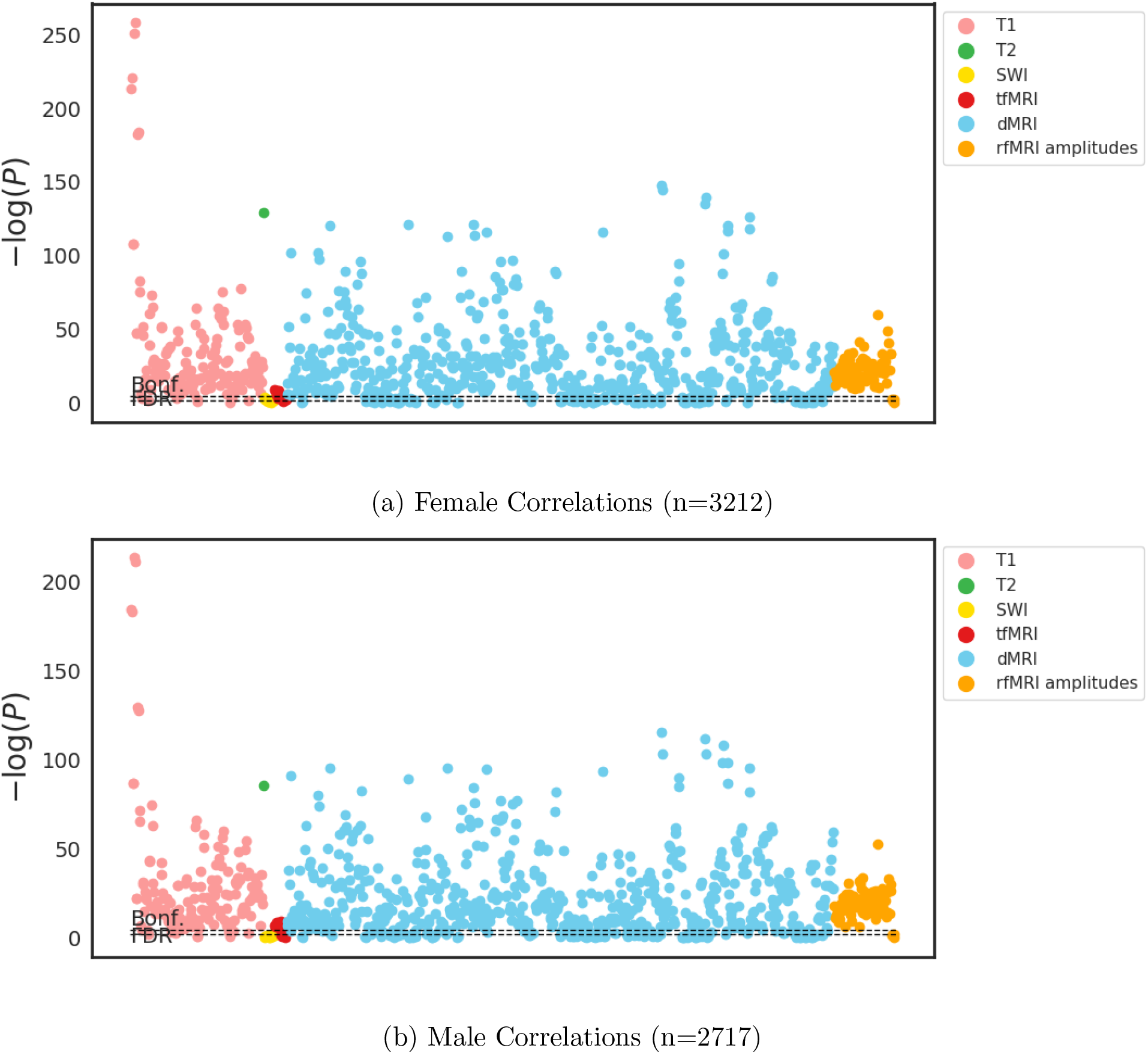
Manhattan plot relating each of the IDPs with the delta measurements from the CNN (with linearly registered images as input) on female and male subjects for the IDP-based method. Each dot represents the statistical significance of the correlation between one IDP type and the deltas. A total of 941 were tested, of which 757 and 751 passed the Bonferroni threshold for the female and male groups, respectively.

**Figure 12:**
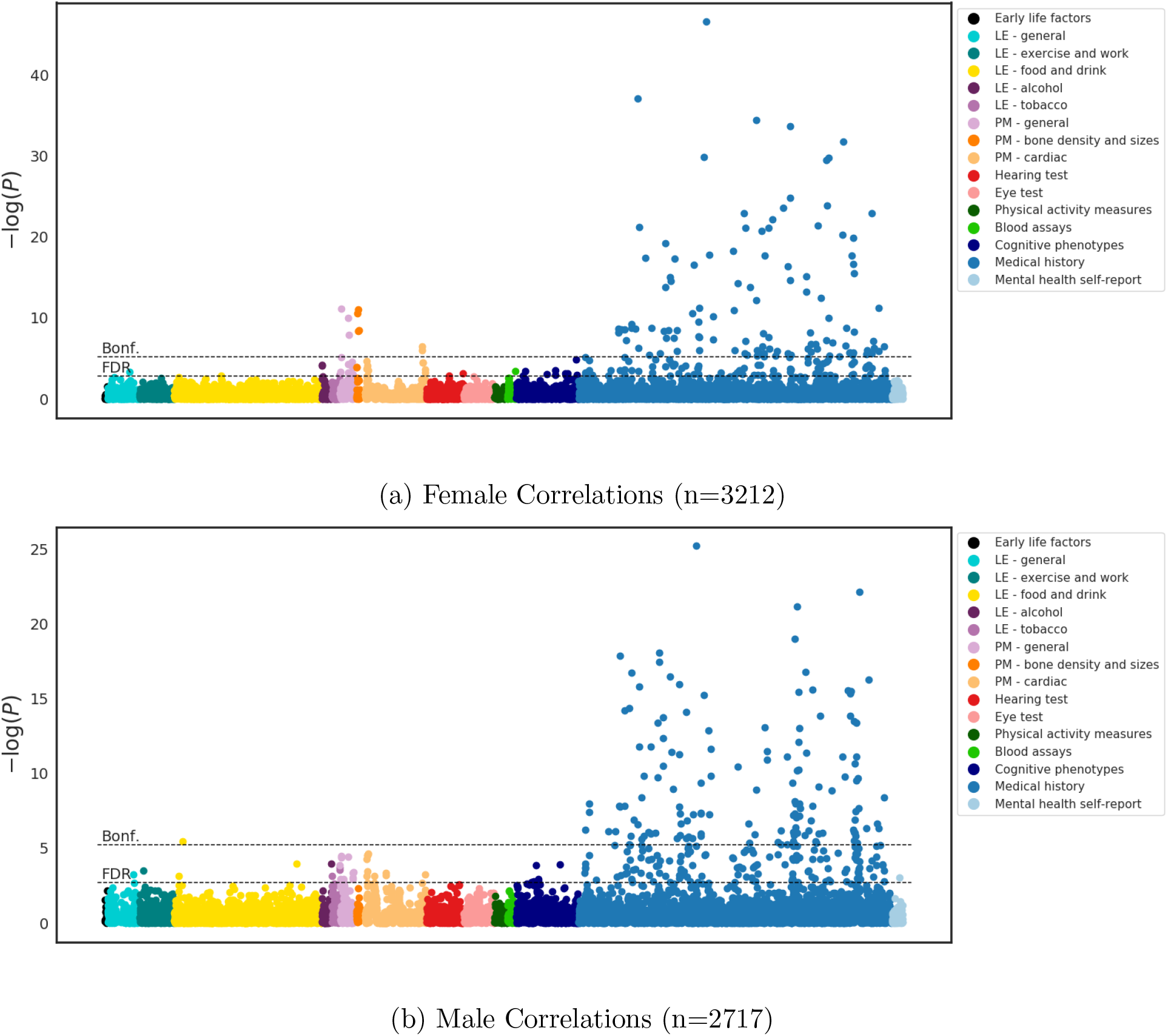
Manhattan plot relating each of the nonimage derived variables with the delta measurements from the CNN (with linearly registered images as input) on female and male subjects for the IDP-based method. Each dot represents the statistical significance of the correlation between one variable type and the deltas. A total of 8787 were tested, of which 120 and 137 passed the Bonferroni threshold for the female and male groups, respectively.

There were biological variables with significant correlations in common between the IDP based method [39] and the CNN predictions. For women, notable common associations were bone density (density, ultrasound attention and mineral density), blood pressure, height, weight, and the presence of MS. For men, the strongest correlations in common were diabetes, hypertension, weight and height. The presence of bone density being a highly correlating factor for women, but not men, follows many reported results from the literature [51]. This was also true when the model was only trained on healthy subjects and tested on all subjects. The common associations between the CNN method and the IDP based method also remained constant.

## 4. Discussion

In this work, a CNN was used to predict brain age using 14, 226 subjects from the UK Biobank as training and a further 6885 subjects for testing. The network performed well on both sex groups, using either the linearly registered or nonlinearly warped images as input. Using the linearly registered image dataset as input, the 3D ensemble network achieved a MAE – for an age range of 44.6 to 80.6 years – of 2.86 2.22 and 3.09 2.37 on the female and male groups respectively (*r* = 0.870 and 0.860). On the nonlinearly warped data the network achieved a MAE of 2.71 2.10 and 2.91 2.18 on the female and male groups respectively (*r* = 0.887 and 0.882). Both of these results are competitive with the MAE range of the 3-5 years achieved for other methods reported in the literature. Although the MAE and MSE scores are lower than most in the current literature, it should be observed that achieving the lowest MAE on the UK Biobank dataset would not yield the most clinically relevant model since not all the subjects are healthy. Subjects 1 and 2 in Fig. 5 have the same true age but very different levels of atrophy and predicted ages. If both subjects were predicted to be the true age, this would result in a lower MAE but would not highlight the large differences in anatomy between the two subjects. This example is striking due to the large structural differences. However, it is entirely possible that there are subjects with pathologies not visible in the T1 images, that the network fails to account for.

The clinical utility of the model could potentially be improved by including information from more imaging modalities. Combining modalities would potentially be better at capturing different aspects of the ageing brain such as brain atrophy, iron deposition, and alterations in white matter microstructure [10, 21] and so improve the model. It is important to note that the ageing process presents in multiple ways: mineralization, microstructure damage, anisotropy loss, and iron deposition [10], and that a T1-weighted MRI is unable to capture all of this information. Despite this, the CNN trained on only T1-weighted images outperformed the multi-modality IDP-based model. Furthermore, as we trained the network using only the T1-weighted MRI, we are able to compute independent correlations with the IDPs from the other modalities. Our model only showed mild correlation with the IDP-based model, however this is not suprising as the IDP-based method is a linear model based on IDPs (i.e., a reduction of voxelwise image data to summary imaging-derived phenotypes) from multiple modalities, whereas our model does not have explicitly defined spatial priors or target structures and, rather, searches the whole image space during the training process. The degree of correlation does indicate a broad agreement between the two methods and the two methods together provide complementary analyses of the data. An interesting further line of investigation could be to explore whether these differences are purely random, driven by noise and non-biological factors, or whether interesting structure exists in the differences, related to different modalities and degrees of nonlinearity.

It is important to consider not only the modality of the input image but the preprocessing applied: in this work, using the non-linearly registered image as input performed as well as using the linearly registered images (*p* values from paired *t*-test between the results from the linearly registered images and nonlinearly registered images: female *p* = 0.710, male *p* = 0.810). This seems surprising because the step of non-linear warping essentially suppresses differences by making the brain conform to the standard template. Whilst certain details and pathologies are still preserved, it is evident in Figs. 5 C and D that the ventricles in the non-linearly registered slices do not appear as dramatically different as they do in the linearly registered slices. This is notable, as change in cortical volume is one of the most common effects of age reported, and many of the methods mentioned have cortical volume or thickness measurements as input for successful age prediction [19, 25, 33]. To roughly examine which brain features the network predicts to be older, all subjects which were predicted to be aged 75-80 by the CNN in each sex group were averaged to form a composite image. This process was repeated for all subjects predicted to be aged 45-50, and the resulting images are shown in Fig. 6. The differences between the 75-80 and 45-50 age groups on the averaged non-linearly registered images are subtle (Fig. 6B); however, the composite image of the linearly registered images of those same subjects (Figure 6A) shows very obvious differences between the age groups. As in Fig. 5, the dramatic differences are most evident in the linearly registered images.

Despite the visual loss of information, the network trained with nonlinearly registered images was able to predict age, and so volumetric information is probably still encoded in the images. Although the nonlinear registration has significantly warped the ventricles, it is still possible to see relatively clear differences in the caudate nucleus, even in the non-linearly registered composite (Fig. 6B). It is possible that this is true for other structures affected by ageing and therefore used by the CNN as features.

As a means for further investigation, attention gates [52] were placed before the first, second, and third max pooling layers, fed with the final layer output, training with both the nonlinearly and linearly registered images Attention gates enhance regions of the image which are useful for the task and suppress the other regions. Fig. 13 shows the output of the attention gates. For the nonlinear case, it indicates that the network is being driven by the areas where the nonlinear registration has had to expand the brain tissue and contract the CSF in the ventricles to match the template (C2). The first gate at the highest resolution locates the brain, and the third gate found few regions of interest at that resolution. More interestingly, C2 was looking at the region around the ventricles, which most likely contains information about how much warping was required to register the ventricles to the MNI template. It can conversely be seen that when training with the linearly registered images, the network considers the brain more as a whole. This would indicate that the network trained on linearly registered images would be more likely to notice subtle changes located away from the ventricles and it is possible that the network trained on nonlinearly registered images might miss cortical changes, which would limit the utility of the network in detecting the early stages of neurological disease. The network used for this can be seen in Fig. 14 in the Supplementary Material. For confirmation and comparison, saliency maps [53] were produced from the network trained without attention gates. In agreement with the attention networks, the saliency maps also indicate that with the nonlinearly registered images, the network focuses only on the area around the ventricles, whereas with the linearly registered images, far more of the image influences the predictions.

**Figure 13:**
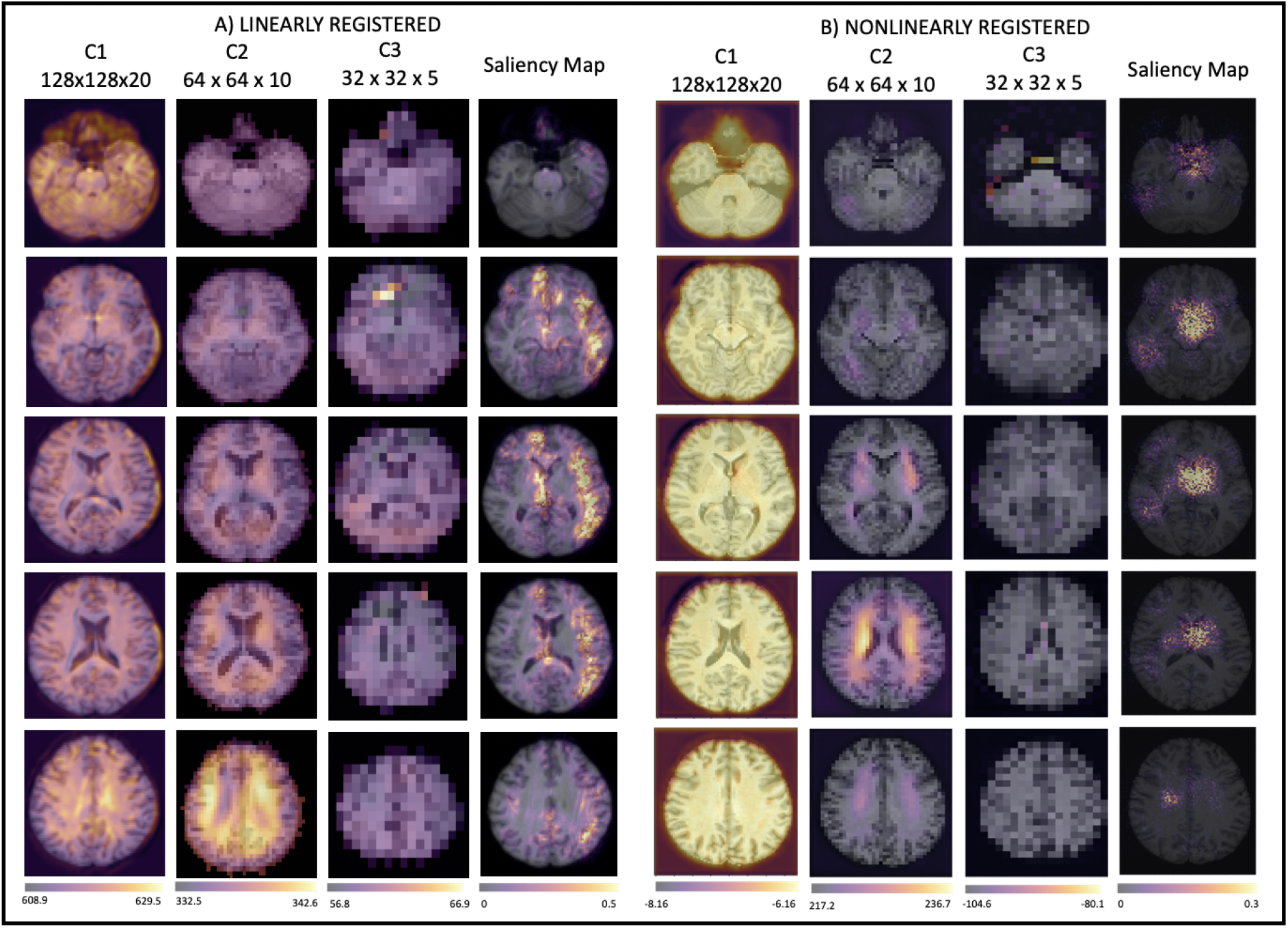
Attention and saliency maps for the two registration types, averaged across all test subjects (n=6885). The attention maps are from three resolutions as indicated by Fig. 14. The saliency maps were produced from the network trained without attention gates.

**Figure 14:**
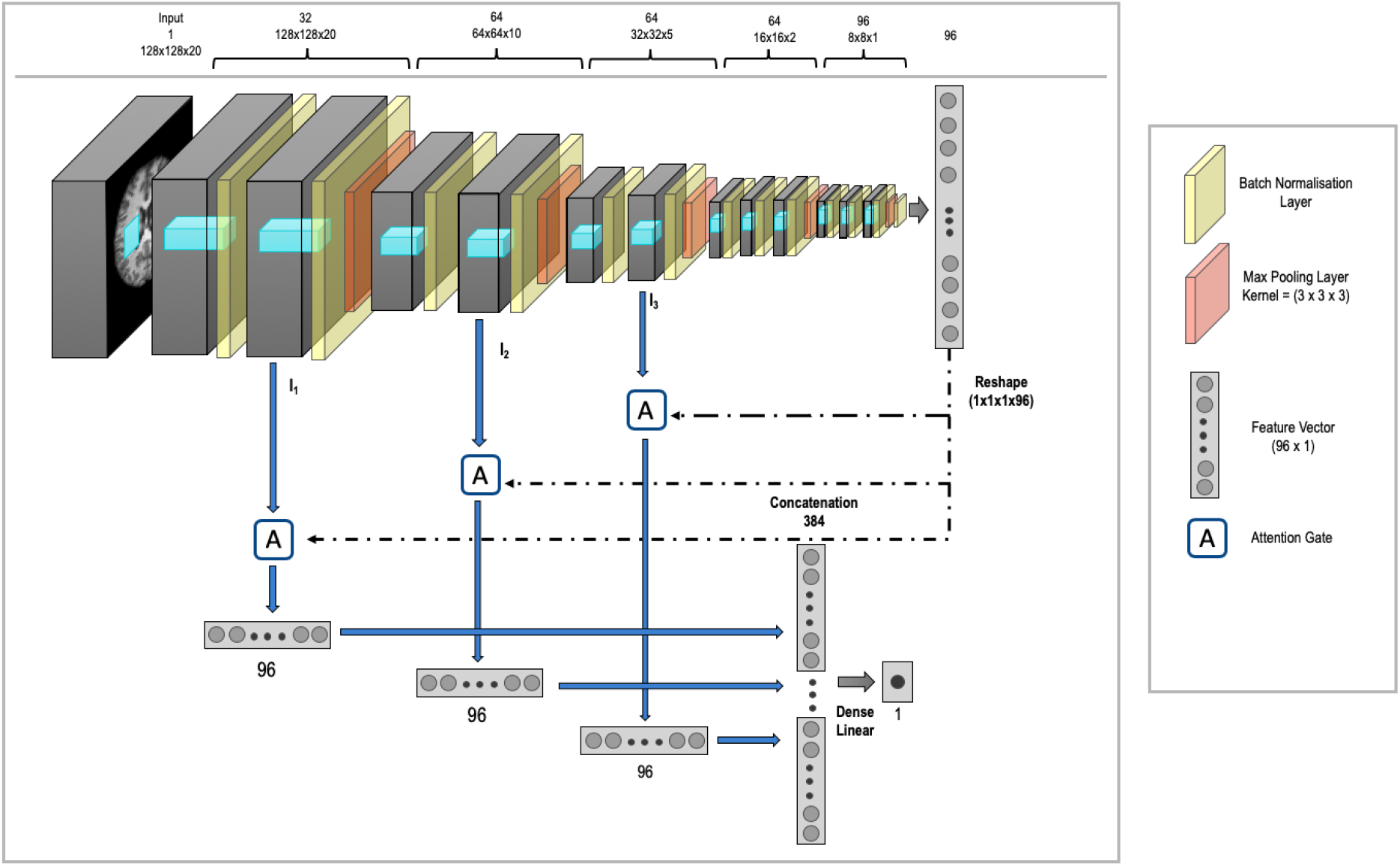
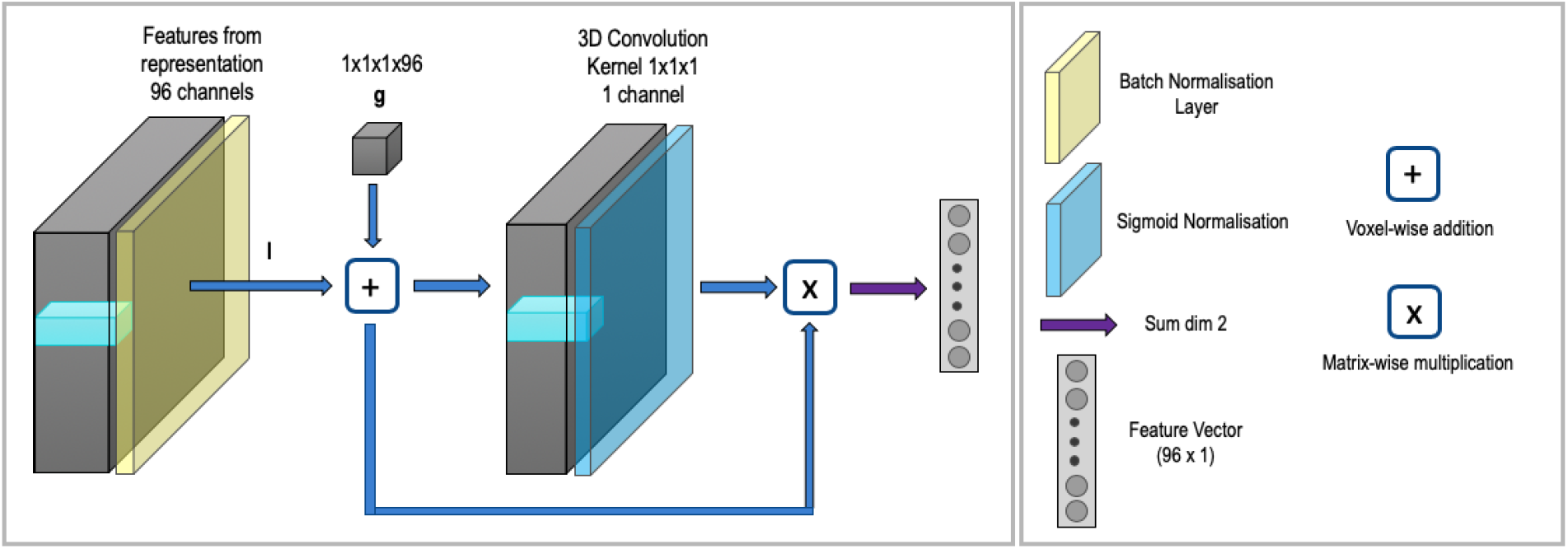
Network used to create the attention maps for the nonlinearly registered data. (a) Network with attention gates added. The spine of the network is the same as is used throughout this work. (b) Attention gate used. Where necessary, the features were first passed through a projection block to ensure the number of features corresponded.

Interestingly, although our network was only trained using only T1 images, the deltas showed significant correlations with 596 and 622 IDPs (in the female and male groups, respectively) from dMRI, T1, and T2 FLAIR images (Figs. 8 and 9). After viewing the anatomical locations of the Tract-Based Spatial Statistics and Probabilistic Tractography IDPs that correlated with the deltas, it was evident that the correlations were distributed throughout the brain and were not particularly localized to a certain region. Overall, one very consistent finding was that intracellular volume fraction (ICVF) showed some of the strongest correlations with the deltas, and the association was always negative. ICVF is related to neuronal density, which means that a higher predicted age than true age (i.e. positive delta) correlated with decreased neuronal density. In addition, IDPs relating to T1 grey matter and white matter volumes correlated negatively with delta, while CSF and the volume of white matter hyperintensities (WMH) correlated positively. This would be expected, since CSF volume in the brain will either increase with age, disease, or loss of other brain tissue, and there is a well-known association between WMH and ageing [54]. The CNN-derived deltas and IDP-derived deltas correlated with mostly similar categories of the UK Biobank variables. In the female subject group, both methods showed correlations passing the Bonferroni threshold with UK Biobank medical measurements from the Physical General, Bone Density and Size, Medical History and Cardiac Physical Measurements categories, as defined by [18]. For the male group, both methods correlated with Medical History and Early Lifestyle (Food and Drink). Only the CNN-based model correlated with the Physical General Measurements. Overall, the variables that correlate with deltas from the CNN are consistent with current knowledge about ageing, and encourage further exploration into the use of brain age as an indicator of healthy or unhealthy ageing. Furthermore, the associations were consistent between training on all subjects or just training on healthy subjects.

The presented method has used one of the greatest number of subjects for CNN-based age prediction, producing accurate age predictions, the errors from which have been to shown to correlate with nonimaging variables suggesting their biological meaningfulness. Already, these findings support the idea that deep learning methods have the potential to generate clinically relevant measurements directly from the medical images themselves. It might be that the errors, or age prediction deltas we are measuring, are useful health predictors, but most subjects do not have diagnosed illnesses. Fortunately, due to the longitudinal aspect of the UK Biobank study, in the future we will have the ability to explore whether the deltas from models such as this were predictive of any health outcomes.

## 5. Data and Code Availability Statement

The data is available from the UK Biobank by application. The code and weights are available from the authors on request through emailing the corresponding author.

## 6. Acknowledgements

This work was supported in parts by the Engineering and Physical Sciences Research Council (EPSRC) and Medical Research Council (MRC) [grant number EP/L016052/1] (N.D, E.B. and Z.A.), the Clarendon fund (E.B), and a Wellcome Trust Strategic Award (098369/Z/12/Z) (S.S. and D.V.). A.N. is grateful for support from the UK Royal Academy of Engineering under the Engineering for Development Research Fellowships scheme. M.J. is supported by the National Institute for Health Research (NIHR) and the Oxford Biomedical Research Centre (BRC). The Wellcome Centre for Integrative Neuroimaging is supported by core funding from the Wellcome Trust (203139/Z/16/Z).

This research has been conducted in part using the UK Biobank Resource under Application Number 8107. We are grateful to UK Biobank for making the data available, and to all UK Biobank study participants, who generously donated their time to make this resource possible. Analysis was carried out on the clusters at the Oxford Biomedical Research Computing (BMRC) facility and FMRIB (part of the Wellcome Centre for Integrative Neuroimaging). BMRC is a joint development between the Wellcome Centre for Human Genetics and the Big Data Institute, supported by Health Data Research UK and the NIHR Oxford Biomedical Research Centre.

Computation used the Oxford Biomedical Research Computing (BMRC) facility, a joint development between the Wellcome Centre for Human Genetics and the Big Data Institute supported by Health Data Research UK and the NIHR Oxford Biomedical Research Centre. The views expressed are those of the author(s) and not necessarily those of the NHS, the NIHR or the Department of Health.

## 7. Supplementary Materials

### 7.1. Age Predictions Results

The age range of the test set for all cases 44.6 - 80.6 years.

### 7.2. Removal of Brain Age Biases

Most brain age literature shows an underestimation of brain age for old subjects and an over estimation of brain age in young subjects, resulting from several factors. This results in an age dependence in the estimated deltas, which will be problematic when computing association with nonimaging variables, as they will be driven by the true age rather than the age delta. We therefore follow the process described by Smith et al [39] to correct for both linear and quadratic associations with age.

We can describe our results in the form:

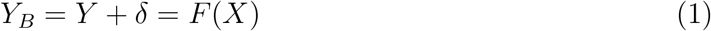

where *Y* are the true age values, *Y*_*B*_ the estimated ‘biased’ age values, *X* the array of brain imaging measurements, which in our case take the form of the T1-weighted MRI images, and F(.) the model mapping from *X* to *Y*_*B*_, which in our case takes the form of the CNN discussed in section 2.2.

We first consider only linear dependence with age, which is simply removed with a second step. If we consider the scatter plots such as those shown in Fig. 4 before the removal of the age dependence, then we can see that ideally we would want the deltas to be distributed around 0 with no overall slope and so no overall dependence on true age Y. If there is the presence of an overall slope then we can simply fit a straight line to the full dataset and subtract this from deltas. Therefore:

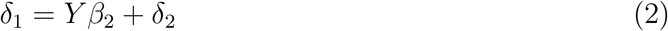

where *β*_2_ is the regressor to model the linear dependence and *δ*_2_ are the residuals from this fitting, which are orthogonal to age with the biases removed. Therefore, the predicted brain age for this step is now:

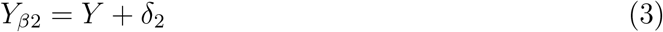

This therefore provides us with an estimate for the deltas with linear relationships to age removed.

This now needs to be extended to include nonlinear relationships with age, especially because the acceleration of the effects of ageing in older age seems likely, especially with disease. We therefore simply adapt the model to include an additive nonlinear term in Y. We simply add a quadratic term as the most natural extension and so:

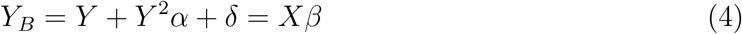

and

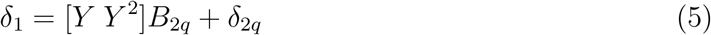

where 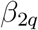 has two regressors covering the linear and quadratic regressors. Therefore, the linear and quadratic relationships with age from the deltas can be removed through multiple regression.

### 7.3. Attention Network Design

### 7.4. Training on Healthy Subjects

As not all of the subjects in the UK Biobank are healthy, we also trained on only the healthy subjects to see what effect this had on the predictions. We did not remove all patients with reported ICD9/10 codes or self-reported conditions as this left very few subjects. Rather, following [55] we removed subjects with conditions that were likely to impact on the brain. We filtered out subjects who had either ICD9/10 codes or self-reported conditions which were likely to influence the brain. Subjects were removed who reported: brain tumours, benign neoplasms of the brain, hydrocephalus/congenital malformations of the nervous system, stroke/cerebrovascular disease, all neurological diagnoses and all mental and behavioural disorders, leaving around 14k subjects in total. The same approach was then taken as explored in the methods but only for the nonlinearly registered images, as this was sufficient for comparison.

The MAE results and correlations can be seen below – Table 5 and Figs. 15 and 16.

**Table 5:**
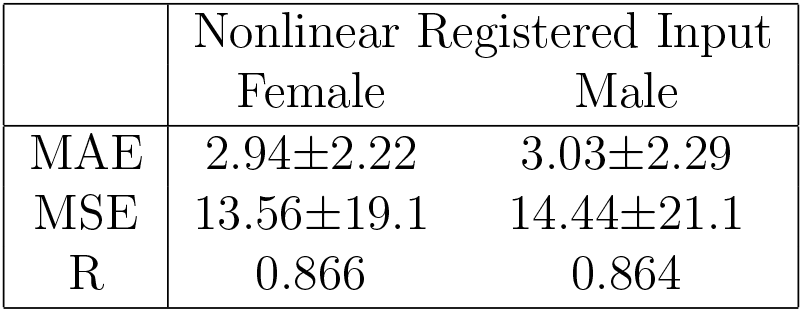
Ensemble Network results split by sex for just the healthy subjects.

**Figure 15:**
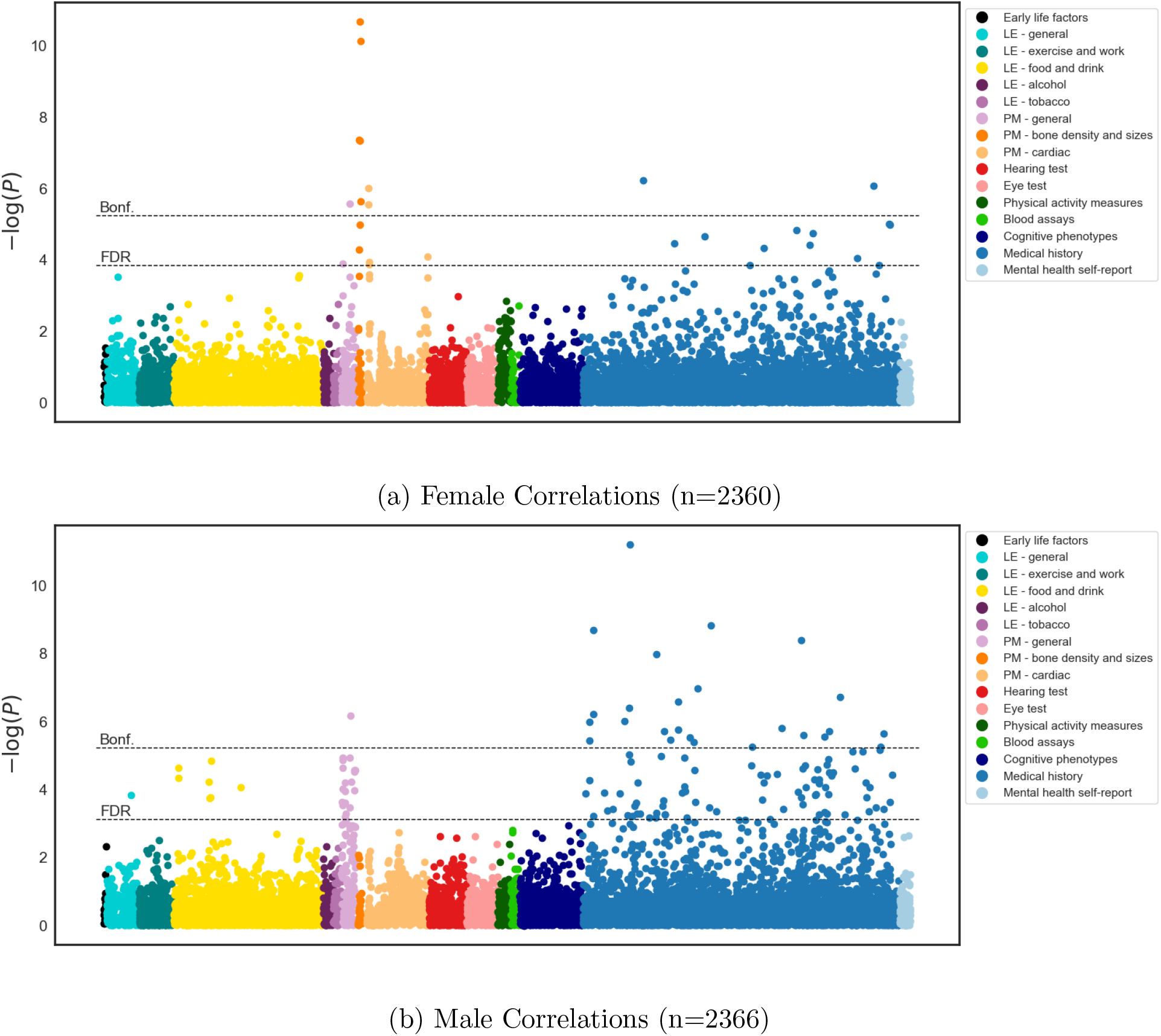
Manhattan plot relating each of the non-imaging UK Biobank variables with the delta measurements from the CNN (with nonlinearly registered images as input) on female and male subjects, training and testing on only the healthy subjects. Each dot represents the statistical significance of the correlation between one UK Biobank variable and the deltas.

**Figure 16:**
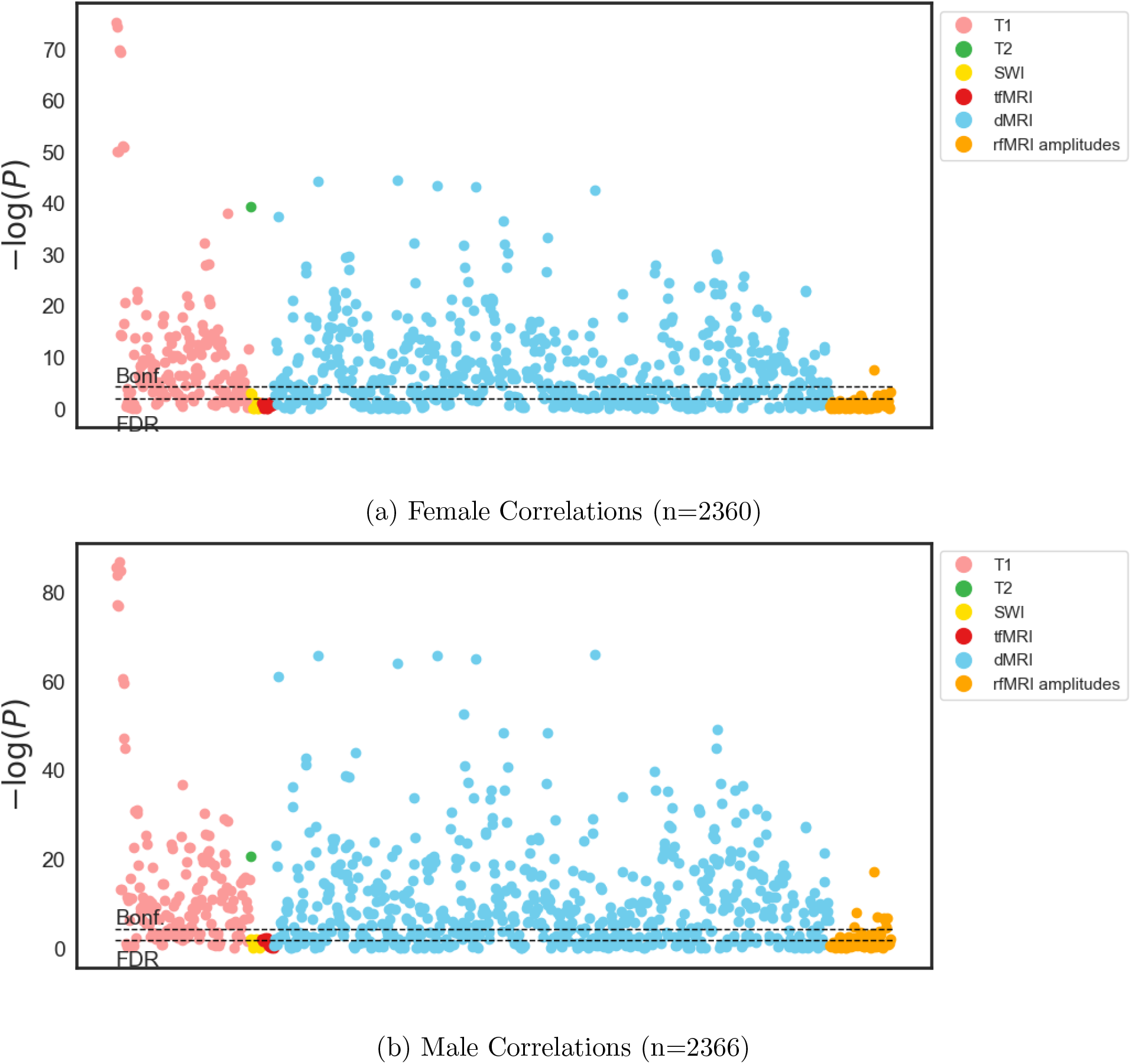
Manhattan plot relating each of the IDPs with the delta measurements from the CNN (with nonlinearly registered images as input) on female and male subjects, training and testing on only the healthy subjects. Each dot represents the statistical significance of the correlation between one UK Biobank variable and the deltas.

We also explored the effect of also removing conditions that have been identified by other works in the literature [39] and by our method to correlate with the Δ_*brainage*_ values – diabetes and hypertension. The removal of these subjects left around 10k subjects in total. Again, the same approach was taken as explored in the methods, but the results are only shown of the nonlinearly registered images as a comparison. The MAE results and correlations are shown – Table 6 and Figs. 17 and 18.

**Table 6:** Ensemble Network results split by sex for just the healthy subjects with correlating conditions removed.

|  | Nonlinear Registered Input |  |
| --- | --- | --- |
|  | Female | Male |
| MAE | 3.20±2.43 | 3.48±2.60 |
| MSE | 16.14±22.9 | 18.91±25.67 |
| R | 0.840 | 0.833 |

**Figure 17:**
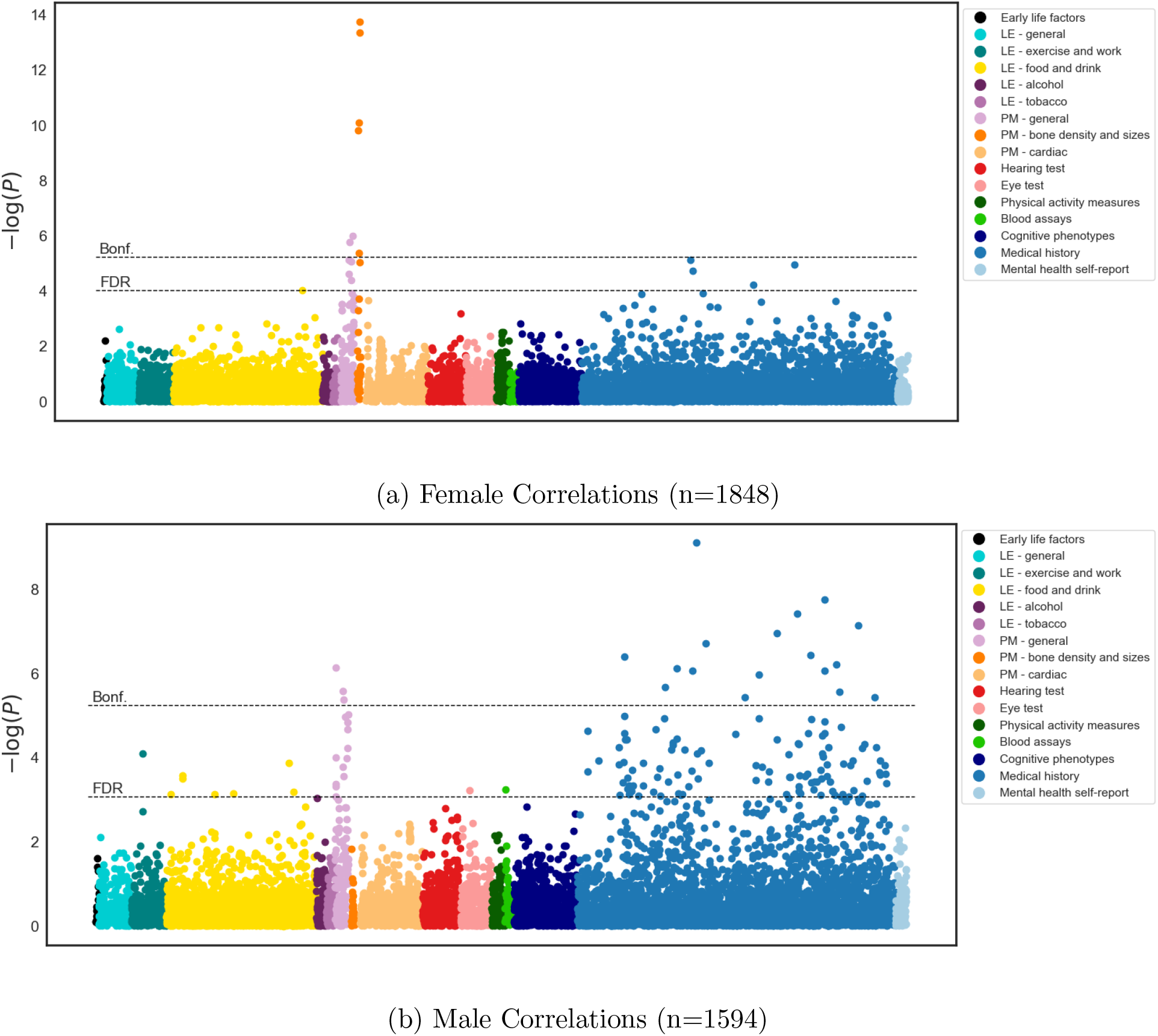
Manhattan plot relating each of the non-imaging UK Biobank variables with the delta measurements from the CNN (with nonlinearly registered images as input) on female and male subjects training and testing on only the healthy subjects, with correlating conditions removed. Each dot represents the statistical significance of the correlation between one UK Biobank variable and the deltas.

**Figure 18:**
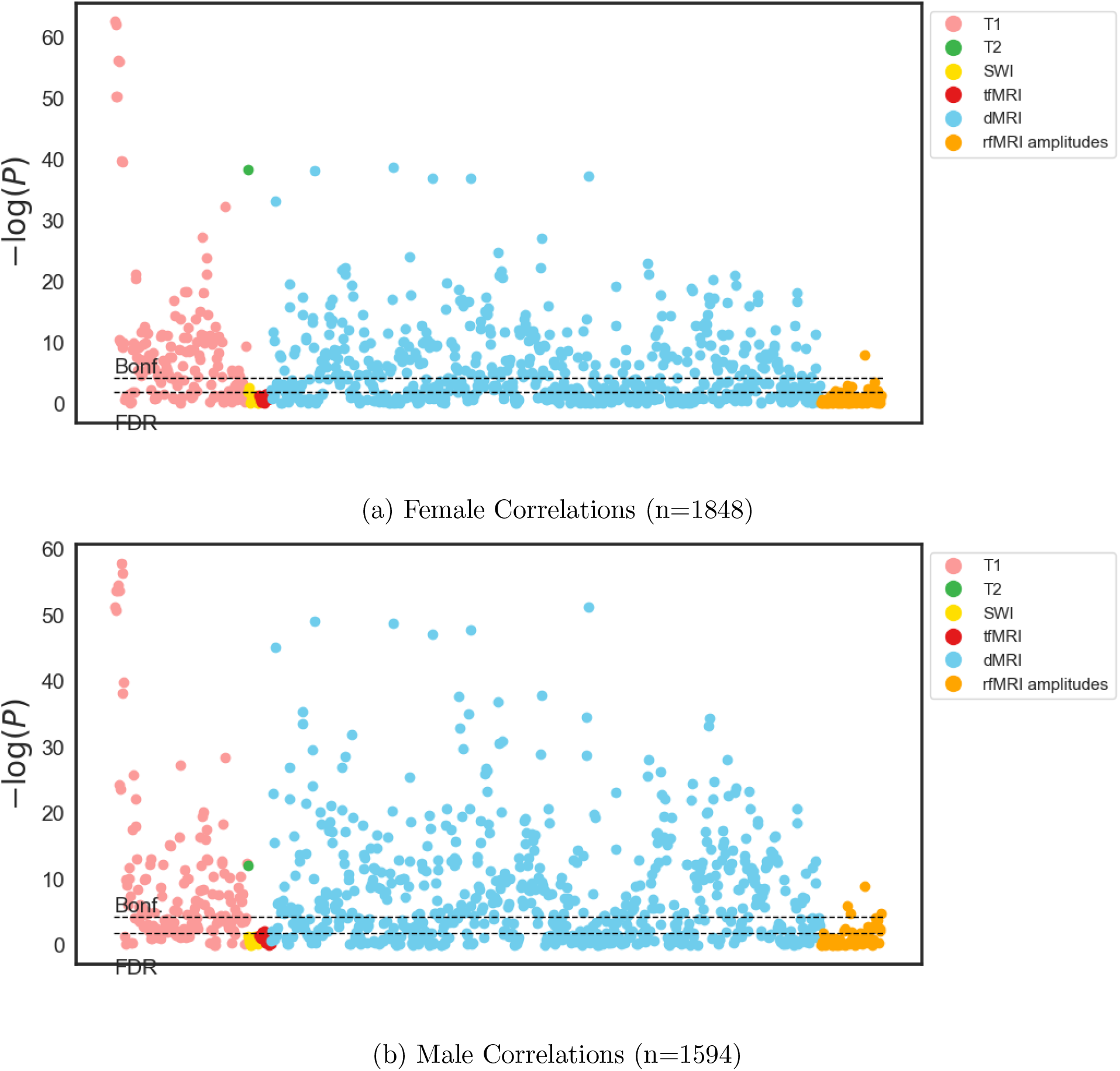
Manhattan plot relating each of the IDPs with the delta measurements from the CNN (with nonlinearly registered images as input) on female and male subjects training and testing on only the healthy subjects, with correlating conditions removed. Each dot represents the statistical significance of the correlation between one UK Biobank variable and the deltas.

Finally, we tested on the whole test dataset – healthy and unhealthy – using the model trained on the healthy subjects, so that subjects with correlating conditions were also removed. The results can be seen in Figs. 19 and 20. It can be seen that the results remain consistent. The highest correlations for women still include bone density, blood pressure, height, weight and the presence of MS, and for men diabetes, hypertension, weight and height.

**Figure 19:**
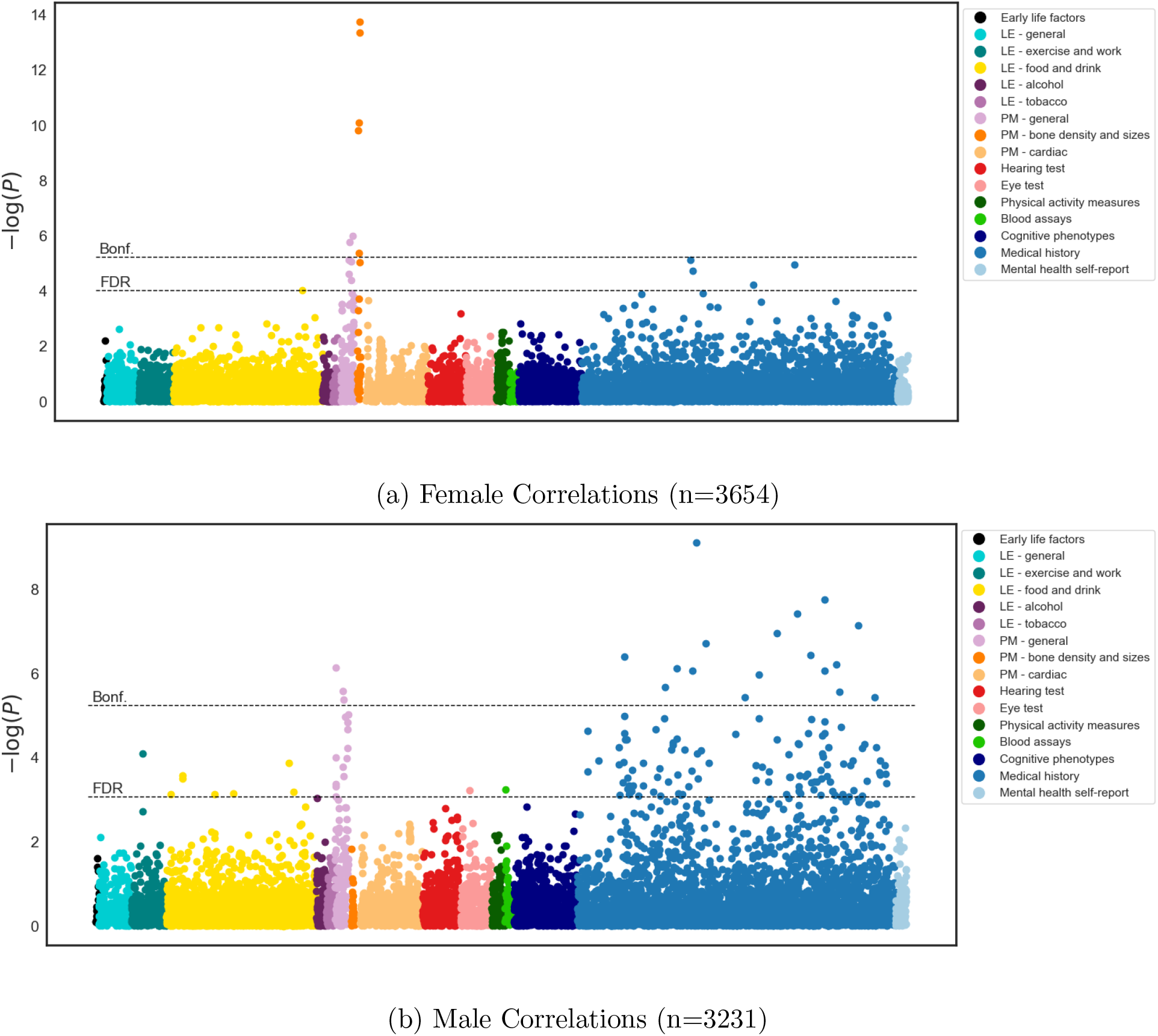
Manhattan plot relating each of the non-imaging UK Biobank variables with the delta measurements from the CNN (with nonlinearly registered images as input) on female and male subjects training on only the healthy subjects, with correlating conditions removed and testing on all subjects. Each dot represents the statistical significance of the correlation between one UK Biobank variable and the deltas.

**Figure 20:**
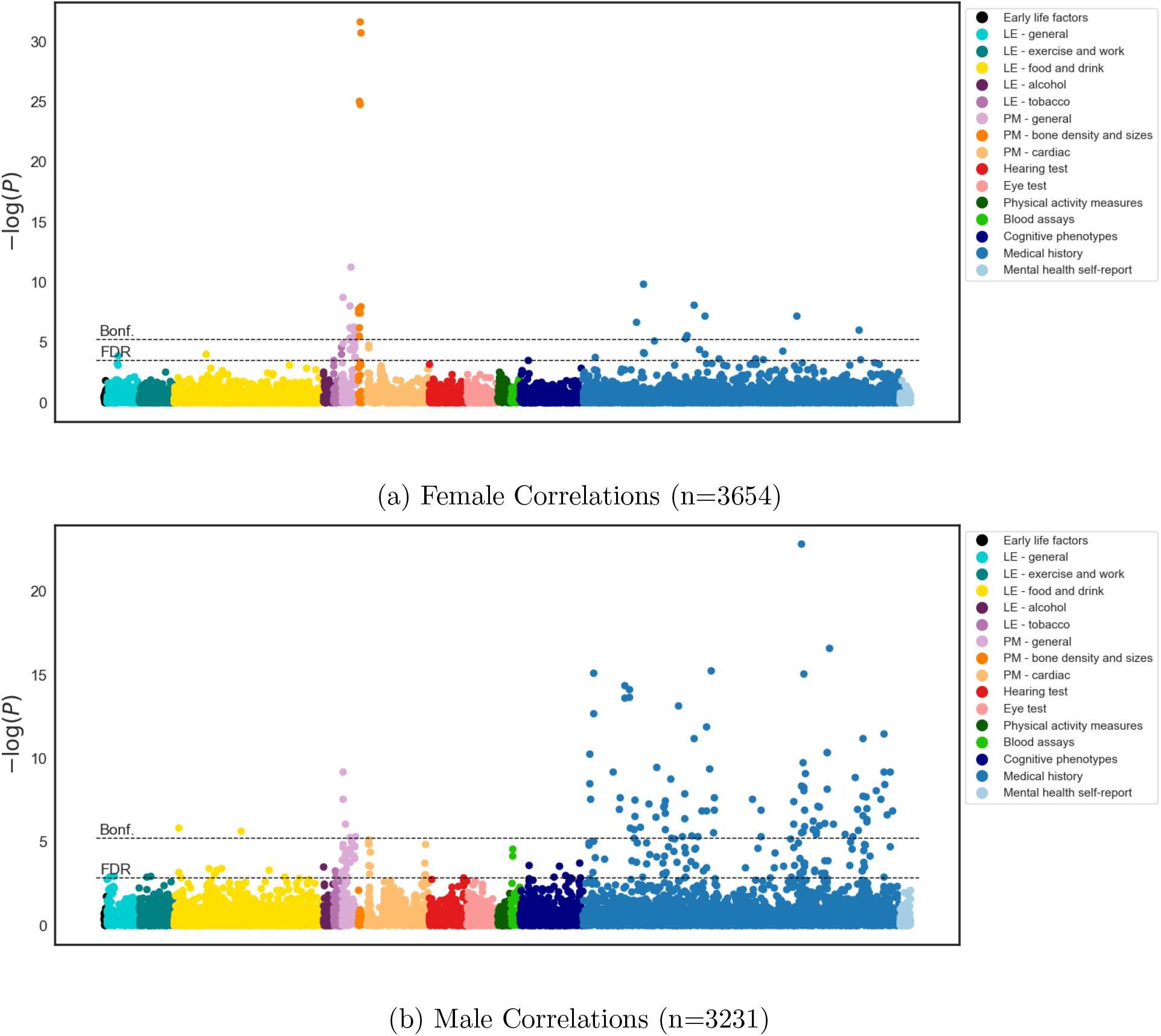
Manhattan plot relating each of the IDPs with the delta measurements from the CNN (with non-linearly registered images as input) on female and male subjects, training on only the healthy subjects with correlating conditions removed and testing on all subjects. Each dot represents the statistical significance of the correlation between one UK Biobank variable and the deltas.

Figure 21 shows the correlations between the deltas from the models trained on the two different healthy cohorts with the model trained on all subjects, where testing was performed using all subjects. It can be seen that there is a high degree of correlation between the deltas produced by the different models, indicating that incorporating the unhealthy subjects into training has little effect on the model predictions. It can also be seen that, while the degree may differ, the models are consistent with the subjects they predict to have accelerated or reduced ageing, meaning that the deltas should be providing meaningful indications of the progression of ageing. Finally it can be seen that there are no subjects that have a large negative delta with one model and large positive delta with the other.

**Figure 21:**
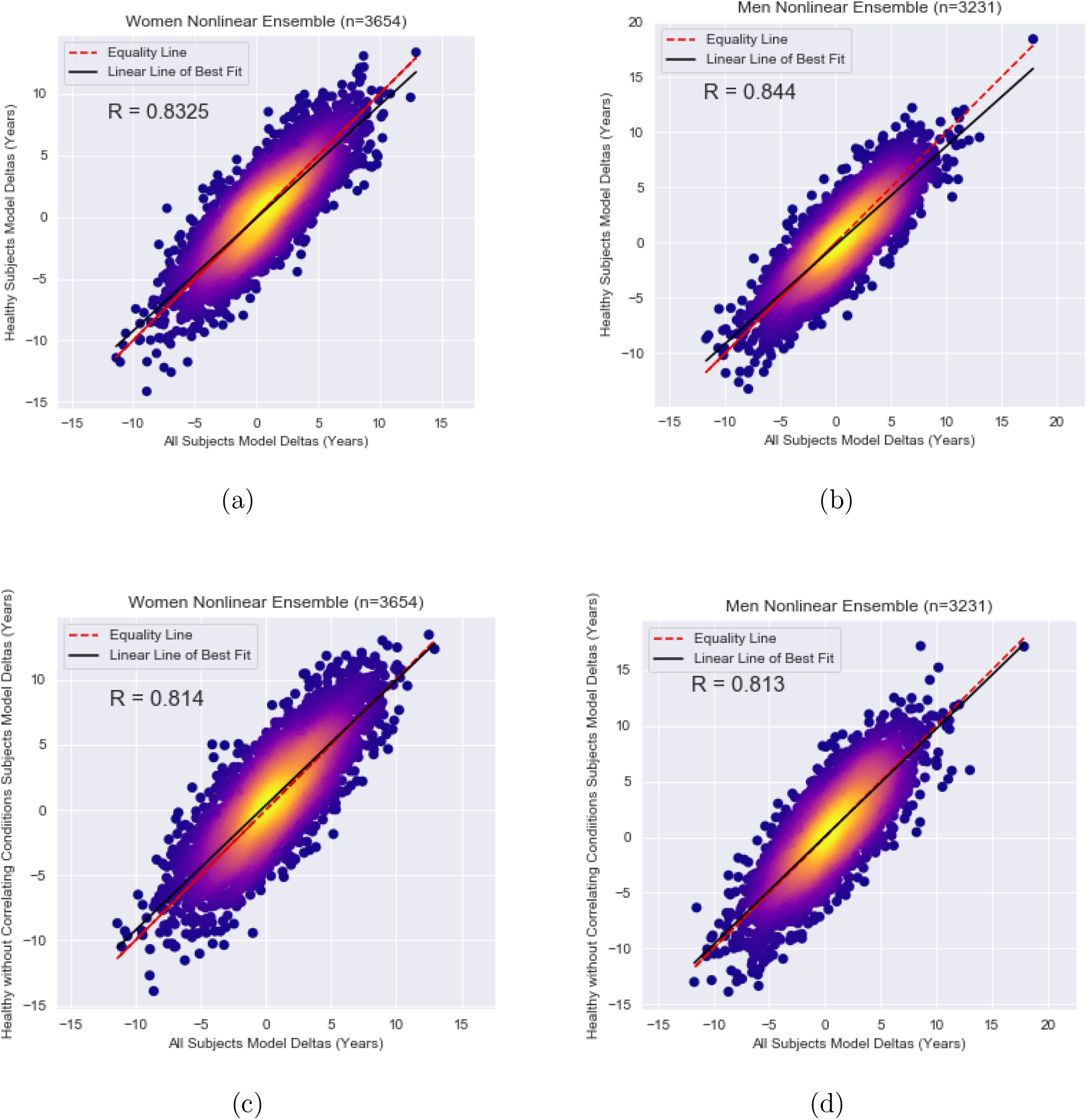
Density plot of the model deltas (true age - pred age) with the model trained on all subjects vs healthy subjects, testing on all subjects (top row) or healthy subjects minus correlating conditions testing on all subjects (bottom row) separated by sex.

### 7.5. Significant IDPS and Variables

For each experiment we present the significant correlations – those which passed Bonferoni correction. As too many IDPS passed the threshold to reasonably to report them all, we show the top 10 for each modality for IDPs and the top 50 for biological variables. The number of the correlation indicates their ranking of significance, with 1 being the most significant.

**Linear Training Data Women:**

**IDPS**:

T1:

1.(r = −0.3), IDP T1 SIENAX grey normalised volume
2.(r = −0.3), IDP T1 SIENAX grey unnormalised volume
3.(r = −0.29), IDP T1 SIENAX peripheral grey normalised volume
4.(r = −0.29), IDP T1 SIENAX peripheral grey unnormalised volume
5.(r = −0.28), IDP T1 SIENAX brain-normalised volume
6.(r = −0.28), IDP T1 SIENAX brain-unnormalised volume
7.(r = 0.27), IDP T1 SIENAX CSF normalised volume
8.(r = 0.27), IDP T1 SIENAX CSF unnormalised volume
14.(r = −0.25), IDP T1 FAST ROIs brain stem
14.(r = −0.25), IDP T1 FAST ROIs brain stem

DMRI:

9.(r = 0.26), IDP dMRI TBSS MD Fornix
10.(r = 0.26), IDP dMRI TBSS L1 Fornix
11.(r = 0.26), IDP dMRI TBSS L2 Fornix
12.(r = 0.26), IDP dMRI TBSS L3 Fornix
13.(r = 0.26), IDP dMRI TBSS ISOVF Fornix
15.(r = −0.25), IDP dMRI TBSS FA Fornix
17.(r = −0.23), IDP dMRI TBSS ICVF Superior fronto-occipital fasciculus L
18.(r = 0.23), IDP dMRI ProbtrackX MD atr r
19.(r = 0.23), IDP dMRI TBSS L2 Fornix cres+Stria terminalis R
20.(r = 0.23), IDP dMRI TBSS L3 Fornix cres+Stria terminalis R

T2:

16.(r = 0.24), IDP T2 FLAIR BIANCA WMH volume

rfMRI:

353.(r = −0.1), rfMRI amplitudes (ICA100 node 38)

**Linear Training Data Women:**

**Non Imaging Variables**:

1. (r =−0.12) [‘Heel broadband ultrasound attenuation (right) (2.0)’]
2. (r =−0.1) [‘Treatment/medication code (1140926934 – tamsulosin)’]
3. (r =−0.1) [‘External causes (W319 – W31.9 Unspecified place)’]
4. (r =−0.1) [‘External causes (W319 – W31.9 Unspecified place)’]
5. (r =−0.1) [‘Heel bone mineral density (BMD) T-score, automated (right) (2.0)’]
6. (r =−0.1) [‘Heel broadband ultrasound attenuation (left) (2.0)’]
7. (r =0.09) [‘Diastolic blood pressure, automated reading (0.1)’]
8. (r =−0.09) [‘Heel bone mineral density (BMD) T-score, automated (left) (2.0)’]
9. (r =−0.08) [‘Weight (2.0)’]
10. (r =0.08) [‘Diagnoses – main ICD10 (G35 – G35 Multiple sclerosis)’]
11. (r =−0.08) [‘Sitting height (2.0)’]
12. (r =−0.08) [‘Sitting height (0.0)’]
13. (r =0.08) [‘Diastolic blood pressure, automated reading (0.0)’]
14. (r =−0.08) [‘Hip circumference (2.0)’]

**Linear Training Data Men:**

**IDPS**:

T1:

1.(r = −0.37), IDP T1 SIENAX grey normalised volume
2.(r = −0.37), IDP T1 SIENAX grey unnormalised volume
3.(r = −0.36), IDP T1 SIENAX peripheral grey normalised volume
4.(r = −0.36), IDP T1 SIENAX peripheral grey unnormalised volume
5.(r = 0.34), IDP T1 SIENAX CSF normalised volume
6.(r = 0.34), IDP T1 SIENAX CSF unnormalised volume
7.(r = −0.33), IDP T1 SIENAX brain-normalised volume
8.(r = −0.33), IDP T1 SIENAX brain-unnormalised volume
17.(r = −0.29), IDP T1 FIRST left thalamus volume
23.(r = −0.28), IDP T1 FIRST right thalamus volume

dMRI:

9.(r = 0.33), IDP dMRI ProbtrackX L1 str r
10.(r = 0.31), IDP dMRI TBSS L2 Fornix cres+Stria terminalis R
11.(r = 0.31), IDP dMRI ProbtrackX L1 str l
12.(r = −0.3), IDP dMRI TBSS ICVF Superior fronto-occipital fasciculus L
13.(r = 0.3), IDP dMRI TBSS L3 Fornix cres+Stria terminalis R
14.(r = 0.3), IDP dMRI TBSS ISOVF Fornix
15.(r = −0.3), IDP dMRI TBSS ICVF Superior fronto-occipital fasciculus R
16.(r = 0.3), IDP dMRI TBSS L2 Fornix
18.(r = −0.3), IDP dMRI TBSS FA Fornix
19.(r = 0.29), IDP dMRI TBSS L3 Fornix

T2:

134.(r = 0.2), IDP T2 FLAIR BIANCA WMH volume

rfMRI:

483.(r = −0.11), rfMRI amplitudes (ICA100 node 17)
500.(r = −0.1), rfMRI amplitudes (ICA100 node 55)
508.(r = −0.1), rfMRI amplitudes (ICA100 node 51)
548.(r = −0.09), rfMRI amplitudes (ICA100 node 54)
567.(r = −0.08), rfMRI amplitudes (ICA100 node 43)
573.(r = −0.08), rfMRI amplitudes (ICA100 node 52)
600.(r = −0.08), rfMRI amplitudes (ICA100 node 21)
611.(r = −0.07), rfMRI amplitudes (ICA100 node 15)

**Linear Training Data Men:**

**Non Imaging Variables:**

1.(r =0.13) [‘Diabetes diagnosed by doctor (0.0)’]
2.(r =0.12) [‘Diabetes diagnosed by doctor (2.0)’]
3.(r =0.12) [‘Non-cancer illness code, self-reported (1220 – diabetes)’]
4.(r =0.11) [‘Treatment/medication code (1140884600 – metformin)’]
5.(r =−0.1) [‘Diagnoses – secondary ICD10 (F402 – F40.2 Specific (isolated) phobias)’]
6.(r =0.1) [‘Non-cancer illness code, self-reported (1065 – hypertension)’]
7.(r =0.1) [‘Medication for cholesterol, blood pressure or diabetes (2.0)’]
9.(r =0.1) [‘Diagnoses – main ICD10 (O60 – O60 Preterm delivery)’]
10.(r=0.1) [‘Diagnoses – secondary ICD10 (E119 – E11.9 Without complications)’]
11.(r=−0.1) [‘Non-cancer illness code, self-reported (1221 – gestational diabetes)’]
12.(r =−0.09) [‘Non-cancer illness code, self-reported (1073 – gestational hypertension/pre-eclampsia)’]
13.(r =0.1) [‘Number of treatments/medications taken (2.0)’]
14.(r =−0.09) [‘Diagnoses – main ICD10 (O16 – O16 Unspecified maternal hypertension)’]
15.(r =−0.09) [‘Diagnoses – secondary ICD10 (Z369 – Z36.9 Antenatal screening, unspecified)’]
16.(r =0.09) [‘Diagnoses – secondary ICD10 (O691 – O69.1 Labour and delivery complicated by cord around neck, with compression)’]
17.(r =0.09) [‘Number of treatments/medications taken (0.0)’]
18.(r =−0.09) [‘Hip circumference (0.0)’]
19.(r =−0.09) [‘Standing height (0.0)’]
20.(r =−0.09) [‘Sitting height (2.0)’]
21.(r =0.08) [‘Treatment/medication code (1140868226 – aspirin)’]
22.(r =−0.08) [‘Cereal intake (2.0)’]
23.(r =0.08) [‘Operation code (1453 – splenectomy)’]
24.(r =−0.09) [‘Standing height (2.0)’]
25.(r =0.08) [‘Treatment/medication code (1140856342 – syndol tablet)’]
26.(r =0.08) [‘Number of self-reported non-cancer illnesses (0.0)’]
27.(r =−0.08) [‘Diagnoses – secondary ICD10 (M255 – M25.5 Pain in joint)’]
28.(r =0.08) [‘Non-cancer illness code, self-reported (1223 – type 2 diabetes)’]

**Nonlinear Training Data Women:**

**IDPS:**

T1:

1.(r = −0.33), IDP T1 SIENAX peripheral grey normalised volume
2.(r = −0.33), IDP T1 SIENAX grey normalised volume
3.(r = −0.33), IDP T1 SIENAX peripheral grey unnormalised volume
4.(r = −0.33), IDP T1 SIENAX grey unnormalised volume
6.(r = −0.28), IDP T1 SIENAX brain-normalised volume
7.(r = −0.28), IDP T1 SIENAX brain-unnormalised volume
20.(r = 0.25), IDP T1 SIENAX CSF normalised volume
21.(r = 0.25), IDP T1 SIENAX CSF unnormalised volume
22.(r = −0.24), IDP T1 FAST ROIs brain stem
51.(r = −0.23), IDP T1 FAST ROIs L cent operc cortex

T2:

5.(r = 0.3), IDP T2 FLAIR BIANCA WMH volume
5.(r = 0.3), IDP T2 FLAIR BIANCA WMH volume

dMRI:

8.(r = 0.29), IDP dMRI ProbtrackX MD atr r
9.(r = 0.28), IDP dMRI ProbtrackX L3 atr r
10.(r = 0.27), IDP dMRI ProbtrackX MD atr l
11.(r = 0.27), IDP dMRI TBSS L3 Fornix cres+Stria terminalis R
12.(r = −0.27), IDP dMRI TBSS ICVF Superior fronto-occipital fasciculus L
13.(r = 0.27), IDP dMRI ProbtrackX L3 atr l
14.(r = 0.27), IDP dMRI ProbtrackX L2 atr r
15.(r = 0.26), IDP dMRI ProbtrackX L1 atr r
16.(r = 0.26), IDP dMRI ProbtrackX L2 atr l
17.(r = 0.25), IDP dMRI TBSS L3 Fornix cres+Stria terminalis L

rfMRI:

321.(r = −0.14), rfMRI amplitudes (ICA100 node 38)
511.(r = −0.09), rfMRI amplitudes (ICA100 node 55)
516.(r = −0.09), rfMRI amplitudes (ICA100 node 43)
523.(r = −0.08), rfMRI amplitudes (ICA100 node 47)
547.(r = −0.08), rfMRI amplitudes (ICA100 node 17)
578.(r = −0.07), rfMRI amplitudes (ICA100 node 23)
580.(r = −0.07), rfMRI amplitudes (ICA100 node 44)

SWI:

535.(r = 0.08), IDP SWI T2star right thalamus

**Nonlinear Training Data Women:**

**Non Imaging Variables**

1. (r =−0.16) [‘Heel broadband ultrasound attenuation (right) (2.0)’]
2. (r =−0.14) [‘Heel bone mineral density (BMD) T-score, automated (right) (2.0)’]
3. (r =−0.13) [‘Heel broadband ultrasound attenuation (left) (2.0)’]
4. (r =−0.12) [‘Heel bone mineral density (BMD) T-score, automated (left) (2.0)’]
5. (r =−0.08) [‘Weight (2.0)’]
6. (r =0.08) [‘Treatment/medication code (1140879802 – amlodipine)’]
7. (r =0.08) [‘Diastolic blood pressure, automated reading (0.1)’]
8. (r =−0.08) [‘Sitting height (0.0)’]
9. (r =−0.1) [‘Heel Broadband ultrasound attenuation, direct entry (0.0)’]
10. (r =−0.1) [‘Heel quantitative ultrasound index (QUI), direct entry (0.0)’]

**Nonlinear Training Data Men:**

**IDPS:**

T1:

1.(r = −0.38), IDP T1 SIENAX grey normalised volume
2.(r = −0.38), IDP T1 SIENAX grey unnormalised volume
3.(r = −0.38), IDP T1 SIENAX peripheral grey normalised volume
4.(r = −0.37), IDP T1 SIENAX peripheral grey unnormalised volume
5.(r = 0.33), IDP T1 SIENAX CSF normalised volume
6.(r = 0.33), IDP T1 SIENAX CSF unnormalised volume
7.(r = −0.33), IDP T1 SIENAX brain-normalised volume
8.(r = −0.33), IDP T1 SIENAX brain-unnormalised volume
34.(r = −0.27), IDP T1 FIRST left thalamus volume
38.(r = −0.27), IDP T1 FIRST right thalamus volume

dMRI:

9.(r = 0.32), IDP dMRI ProbtrackX L1 str l
10.(r = 0.32), IDP dMRI ProbtrackX L1 str r
11.(r = 0.32), IDP dMRI TBSS L2 Fornix cres+Stria terminalis R
12.(r = −0.32), IDP dMRI TBSS ICVF Superior fronto-occipital fasciculus L
13.(r = 0.31), IDP dMRI TBSS L3 Fornix cres+Stria terminalis R
14.(r = 0.31), IDP dMRI TBSS L2 Fornix
15.(r = 0.31), IDP dMRI TBSS L3 Fornix
16.(r = −0.31), IDP dMRI TBSS FA Fornix
17.(r = 0.31), IDP dMRI TBSS MD Fornix
18.(r = 0.31), IDP dMRI TBSS ISOVF Fornix

T2:

130.(r = 0.21), IDP T2 FLAIR BIANCA WMH volume

rfMRI:

234.(r = −0.17), rfMRI amplitudes (ICA100 node 38)
428.(r = −0.12), rfMRI amplitudes (ICA100 node 17)
484.(r = −0.11), rfMRI amplitudes (ICA100 node 55)
503.(r = −0.11), rfMRI amplitudes (ICA100 node 15)
511.(r = −0.11), rfMRI amplitudes (ICA100 node 43)
522.(r = −0.1), rfMRI amplitudes (ICA100 node 51)
527.(r = −0.1), rfMRI amplitudes (ICA100 node 21)
551.(r = −0.09), rfMRI amplitudes (ICA100 node 54)
566.(r = −0.09), rfMRI amplitudes (ICA100 node 52)
580.(r = −0.08), rfMRI amplitudes (ICA100 node 37)

**Nonlinear Training Data Men:**

**Non Imaging Variables**

1. (r =0.14) [‘Diabetes diagnosed by doctor (2.0)’]
2. (r =0.14) [‘Non-cancer illness code, self-reported (1220 – diabetes)’]
3. (r =0.14) [‘Treatment/medication code (1140884600 – metformin)’]
4. (r =0.12) [‘Diabetes diagnosed by doctor (0.0)’]
5. (r =0.1) [‘Non-cancer illness code, self-reported (1065 – hypertension)’]
6. (r =−0.1) [‘Standing height (0.0)’]
7. (r =0.1) [‘Treatment/medication code (1140874744 – gliclazide)’]
8. (r =−0.11) [‘Standing height (2.0)’]
9. (r =0.1) [‘Treatment/medication code (1140868226 – aspirin)’]
10. (r =0.1) [‘Medication for cholesterol, blood pressure or diabetes (2.0)’]
11. (r =0.11) [‘Number of treatments/medications taken (2.0)’]
12. (r =0.1) [‘Diagnoses – secondary ICD10 (E119 – E11.9 Without complications)’]
13. (r =−0.1) [‘Diagnoses – secondary ICD10 (F402 – F40.2 Specific (isolated) phobias)’]
14. (r =−0.1) [‘Diagnoses – secondary ICD10 (M255 – M25.5 Pain in joint)’]
15. (r =−0.1) [‘Trunk fat-free mass (0.0)’]
16. (r =−0.1) [‘Arm fat-free mass (left) (0.0)’]
17. (r =−0.1) [‘Arm predicted mass (left) (0.0)’]
18. (r =0.09) [‘Diagnoses – main ICD10 (O60 – O60 Preterm delivery)’]
19. (r =−0.1) [‘Sitting height (2.0)’]
20. (r =−0.09) [‘Non-cancer illness code, self-reported (1221 – gestational diabetes)’]
21. (r =−0.09) [‘Basal metabolic rate (0.0)’]
22. (r =0.09) [‘Number of treatments/medications taken (0.0)’]
23. (r =−0.09) [‘Arm predicted mass (right) (0.0)’]
24. (r =−0.09) [‘Diagnoses – main ICD10 (O16 – O16 Unspecified maternal hypertension)’]
25. (r =0.09) [‘Non-cancer illness code, self-reported (1223 – type 2 diabetes)’]
26. (r =−0.09) [‘Weight (2.0)’]
27. (r =0.09) [‘Diagnoses – main ICD10 (O034 – O03.4 Incomplete, without complication)’]
28. (r =0.08) [‘Treatment/medication code (1140860806 – ramipril)’]
29. (r =0.19) [‘Diabetes diagnosed by doctor (1.0)’]
30. (r =0.08) [‘Medication for cholesterol, blood pressure or diabetes (0.0)’]
31. (r =−0.08) [‘Cereal intake (2.0)’]
32. (r =0.08) [‘Treatment/medication code (1140883066 – insulin product)’]
33. (r =0.08) [‘Treatment/medication code (1140879760 – bisoprolol)’]
34. (r =−0.08) [‘Diagnoses – main ICD10 (Z131 – Z13.1 Special screening examination for diabetes mellitus)’]
35. (r =0.08) [‘Non-cancer illness code, self-reported (1470 – anorexia/bulimia/other eating disorder)’]
36. (r =−0.08) [‘Weight (0.0)’]

**IDP Based Method Women:**

**IDPS:**

T1:

1.(r = −0.56), IDP T1 SIENAX grey unnormalised volume
2.(r = −0.55), IDP T1 SIENAX grey normalised volume
3.(r = −0.52), IDP T1 SIENAX peripheral grey unnormalised volume
4.(r = −0.52), IDP T1 SIENAX peripheral grey normalised volume
5.(r = −0.48), IDP T1 SIENAX brain-unnormalised volume
6.(r = −0.48), IDP T1 SIENAX brain-normalised volume
23.(r = 0.38), IDP T1 SIENAX CSF normalised volume
24.(r = 0.38), IDP T1 SIENAX CSF unnormalised volume
44.(r = −0.33), IDP T1 FIRST right thalamus volume
49.(r = −0.32), IDP T1 FAST ROIs brain stem

dMRI:

7.(r = 0.44), IDP dMRI ProbtrackX MD atr l
8.(r = 0.43), IDP dMRI ProbtrackX MD atr r
9.(r = 0.43), IDP dMRI ProbtrackX L1 atr r
10.(r = 0.42), IDP dMRI ProbtrackX L1 atr l
12.(r = 0.41), IDP dMRI ProbtrackX L3 atr l
13.(r = 0.4), IDP dMRI TBSS L2 Fornix cres+Stria terminalis R
14.(r = 0.4), IDP dMRI TBSS L1 Fornix
15.(r = 0.4), IDP dMRI ProbtrackX L2 atr l 16.(r = 0.4), IDP dMRI TBSS MD Fornix
17.(r = 0.4), IDP dMRI ProbtrackX L3 atr r

T2:

11.(r = 0.41), IDP T2 FLAIR BIANCA WMH volume

rfMRI:

91.(r = −0.29), rfMRI amplitudes (ICA100 node 38)
135.(r = −0.26), rfMRI amplitudes (ICA100 node 51)
185.(r = −0.24), rfMRI amplitudes (ICA100 node 15)
191.(r = −0.24), rfMRI amplitudes (ICA100 node 52)
212.(r = −0.23), rfMRI amplitudes (ICA100 node 21)
248.(r = −0.22), rfMRI amplitudes (ICA100 node 8)
253.(r = −0.21), rfMRI amplitudes (ICA100 node 2)
255.(r = −0.21), rfMRI amplitudes (ICA100 node 55)
263.(r = −0.21), rfMRI amplitudes (ICA100 node 44)
265.(r = −0.21), rfMRI amplitudes (ICA100 node 54)
tfMRI: 736.(r = −0.08), IDP tfMRI median zstat shapes
739.(r = −0.08), IDP tfMRI median zstat faces

**IDP Based Method Women:**

**Non Imaging Variables:**

1. (r =0.23) [‘Non-cancer illness code, self-reported (1507 – haemochromatosis)’]
2. (r =0.22) [‘Diagnoses – main ICD10 (E119 – E11.9 Without complications)’]
3. (r =−0.22) [‘Diagnoses – main ICD10 (N433 – N43.3 Hydrocele, unspecified)’]
4. (r =0.21) [‘Diagnoses – secondary ICD10 (M1099 – M10.99 Gout, unspecified (Site unspecified))’]
5. (r =−0.2) [‘Operation code (1453 – splenectomy)’]
6. (r =0.2) [‘Diagnoses – secondary ICD10 (I460 – I46.0 Cardiac arrest with successful resuscitation)’]
7. (r =0.2) [‘Diagnoses – secondary ICD10 (I120 – I12.0 Hypertensive renal disease with renal failure)’]
8. (r =0.18) [‘Diagnoses – main ICD10 (N47 – N47 Redundant prepuce, phimosis and paraphimossis)’]
9. (r =0.18) [‘Diagnoses – secondary ICD10 (I248 – I24.8 Other forms of acute ischaemic heart disease)’]
10. (r =0.18) [‘Diagnoses – main ICD10 (M2551 – M25.51 Pain in joint (Shoulder region))’]
11. (r =0.18) [’External causes (W319 – W31.9 Unspecified place)’]
12. (r =0.18) [‘External causes (W319 – W31.9 Unspecified place)’]
13. (r =0.17) [‘Diagnoses – main ICD10 (K403 – K40.3 Unilateral or unspecified inguinal hernia, with obstruction, without gangrene)’]
14. (r =0.17) [‘Non-cancer illness code, self-reported (1519 – kidney nephropathy)’]
15. (r =−0.17) [‘Diagnoses – main ICD10 (K081 – K08.1 Loss of teeth due to accident, extraction or local periodontal disease)’]
16. (r =0.17) [‘External causes (Y400 – Y40.0 Penicillins)’]
17. (r =−0.17) [‘Diagnoses – main ICD10 (H810 – H81.0 Meniere’s disease)”]
18. (r =0.17) [‘Diagnoses – secondary ICD10 (M109 – M10.9 Gout, unspecified)’]
19. (r =0.16) [‘Treatment/medication code (1140874686 – glucophage 500mg tablet)’]
20. (r =−0.16) [‘Operation code (1526 – stapedectomy)’]
21. (r =−0.15) [‘Diagnoses – main ICD10 (I840 – I84.0 Internal thrombosed haemorrhoids)’]
22. (r =0.15) [‘Diagnoses – secondary ICD10 (N210 – N21.0 Calculus in bladder)’]
23. (r =−0.15) [‘Non-cancer illness code, self-reported (1629 – fracture face / orbit / eye socket)’]
24. (r =0.15) [‘Treatment/medication code (1140888594 – fluvastatin)’]
25. (r =0.15) [‘Treatment/medication code (1141187810 – tadalafil)’]
26. (r =0.15) [‘Diagnoses – main ICD10 (N210 – N21.0 Calculus in bladder)’]
27. (r =0.14) [‘Diagnoses – main ICD10 (S430 – S43.0 Dislocation of shoulder joint)’]
28. (r =0.14) [‘Treatment/medication code (1140881938 – beclomethasone dipropionate+salbutamol)’]
29. (r =0.14) [‘Diagnoses – main ICD10 (N481 – N48.1 Balanoposthitis)’]
30. (r =−0.14) [‘Treatment/medication code (1140882498 – penicillin)’]
31. (r =0.14) [‘Histology of cancer tumour (9061 – Seminoma, NOS)’]
32. (r =0.14) [‘Treatment/medication code (1140875408 – allopurinol)’]
33. (r =−0.13) [‘Diagnoses – main ICD10 (S4200 – S42.00 Fracture of clavicle (closed))’]
34. (r =0.13) [‘Diagnoses – secondary ICD10 (E831 – E83.1 Disorders of iron metabolism)’]
35. (r =0.13) [‘Diagnoses – main ICD10 (F100 – F10.0 Acute intoxication)’]
36. (r =0.12) [‘Diagnoses – secondary ICD10 (Z572 – Z57.2 Occupational exposure to dust)’]
37. (r =−0.13) [‘Hip circumference (2.0)’]
38. (r =−0.13) [‘Heel broadband ultrasound attenuation (right) (2.0)’]
39. (r =−0.12) [‘Type of cancer: ICD10 (D046 – D04.6 Skin of upper limb, including shoulder)’]
40. (r =0.12) [‘Treatment/medication code (1141179974 – cozaar 25mg tablet)’]
41. (r =−0.13) [‘Heel broadband ultrasound attenuation (left) (2.0)’]
42. (r =−0.12) [‘Body mass index (BMI) (2.0)’]
43. (r =0.11) [‘Diagnoses – secondary ICD10 (I420 – I42.0 Dilated cardiomyopathy)’]
44. (r =0.11) [‘Operation code (1108 – leg artery angioplasty +/− stent)’]
45. (r =0.11) [‘Non-cancer illness code, self-reported (1408 – alcohol dependency)’]
46. (r =0.11) [‘Diagnoses – secondary ICD10 (M2551 – M25.51 Pain in joint (Shoulder region))’]
47. (r =0.11) [‘Non-cancer illness code, self-reported (1400 – peptic ulcer)’]
48. (r =0.11) [‘Treatment/medication code (1140860806 – ramipril)’]
49. (r =0.11) [‘Non-cancer illness code, self-reported (1466 – gout)’]
50. (r =0.11) [‘Non-cancer illness code, self-reported (1121 – pulmonary fibrosis)’]

**IDP Based Method Men:**

**IDPS:**

T1:

1.(r = −0.55), IDP T1 SIENAX grey normalised volume
2.(r = −0.55), IDP T1 SIENAX grey unnormalised volume
3.(r = −0.52), IDP T1 SIENAX peripheral grey normalised volume
4.(r = −0.51), IDP T1 SIENAX peripheral grey unnormalised volume
5.(r = −0.44), IDP T1 SIENAX brain-normalised volume
6.(r = −0.44), IDP T1 SIENAX brain-unnormalised volume
23.(r = 0.37), IDP T1 SIENAX CSF normalised volume
24.(r = 0.37), IDP T1 SIENAX CSF unnormalised volume
36.(r = −0.34), IDP T1 FAST ROIs L frontal pole
39.(r = −0.34), IDP T1 FIRST left thalamus volume

dMRI:

7.(r = 0.42), IDP dMRI ProbtrackX MD atr l
8.(r = 0.41), IDP dMRI ProbtrackX L1 atr l
9.(r = 0.41), IDP dMRI ProbtrackX L1 str r
10.(r = 0.4), IDP dMRI ProbtrackX MD atr r
11.(r = 0.4), IDP dMRI ProbtrackX L1 atr r
12.(r = 0.39), IDP dMRI ProbtrackX L1 str l
13.(r = 0.39), IDP dMRI ProbtrackX L2 atr l
14.(r = 0.38), IDP dMRI TBSS MD Fornix
15.(r = 0.38), IDP dMRI TBSS L2 Fornix
16.(r = 0.38), IDP dMRI ProbtrackX L3 atr l

T2:

25.(r = 0.36), IDP T2 FLAIR BIANCA WMH volume

rfMRI:

85.(r = −0.29), rfMRI amplitudes (ICA100 node 38)
199.(r = −0.23), rfMRI amplitudes (ICA100 node 19)
201.(r = −0.23), rfMRI amplitudes (ICA100 node 52)
211.(r = −0.23), rfMRI amplitudes (ICA100 node 15)
229.(r = −0.22), rfMRI amplitudes (ICA100 node 17)
233.(r = −0.22), rfMRI amplitudes (ICA100 node 2)
239.(r = −0.22), rfMRI amplitudes (ICA100 node 54)
265.(r = −0.21), rfMRI amplitudes (ICA100 node 18)
270.(r = −0.21), rfMRI amplitudes (ICA100 node 44)
278.(r = −0.21), rfMRI amplitudes (ICA100 node 47)

tfMRI:

635.(r = −0.12), IDP tfMRI 90th-percentile zstat faces
643.(r = −0.11), IDP tfMRI median zstat shapes
652.(r = −0.11), IDP tfMRI 90th-percentile zstat shapes
657.(r = −0.11), IDP tfMRI median zstat faces
687.(r = −0.1), IDP tfMRI median BOLD shapes
710.(r = −0.1), IDP tfMRI median BOLD faces
718.(r = −0.09), IDP tfMRI 90th-percentile BOLD shapes
723.(r = −0.09), IDP tfMRI 90th-percentile BOLD faces
729.(r = −0.09), IDP tfMRI 90th-percentile zstat faces-shapes

**IDP Based Method Men:**

**Non Imaging Variables:**

1. (r =0.2) [‘Treatment/medication code (1141195224 – formoterol)’]
2. (r =−0.19) [‘Diagnoses – secondary ICD10 (O701 – O70.1 Second degree perineal laceration during delivery)’]
3. (r =0.18) [‘Diagnoses – main ICD10 (O16 – O16 Unspecified maternal hypertension)’]
4. (r =−0.17) [‘Treatment/medication code (1140868460 – progynova 1mg tablet)’]
5. (r =−0.16) [‘Diagnoses – main ICD10 (R935 – R93.5 Abnormal findings on diagnostic imaging of other abdominal regions, including retroperitoneum)’]
6. (r =0.16) [‘Treatment/medication code (1140881320 – magnesium carbonate)’]
7. (r =0.16) [‘Diagnoses – secondary ICD10 (T884 – T88.4 Failed or difficult intubation)’]
8. (r =0.16) [‘Diagnoses – main ICD10 (Z411 – Z41.1 Other plastic surgery for unacceptable cosmetic appearance)’]
9. (r =0.16) [‘Diagnoses – secondary ICD10 (M511 – M51.1 Lumbar and other intervertebral disk disorders with radiculopathy)’]
10. (r =0.16) [‘Diagnoses – secondary ICD10 (M790 – M79.0 Rheumatism, unspecified)’]
11. (r =−0.15) [‘Operation code (1453 – splenectomy)’]
12. (r =0.15) [‘Non-cancer illness code, self-reported (1221 – gestational diabetes)’]
13. (r =0.15) [‘Treatment/medication code (1141157458 – hypromellose product)’]
14. (r =0.15) [‘Diagnoses – secondary ICD10 (M7909 – M79.09 Rheumatism, unspecified (Site unspecified))’]
15. (r =0.15) [‘Treatment/medication code (1140872036 – paramax tablet)’]
16. (r =−0.14) [‘Diagnoses – secondary ICD10 (N701 – N70.1 Chronic salpingitis and oophoritis)’]
17. (r =0.14) [‘Treatment/medication code (1140866280 – bumetanide)’]
18. (r =−0.14) [‘Diagnoses – main ICD10 (I843 – I84.3 External thrombosed haemorrhoids)’]
19. (r =−0.14) [‘Diagnoses – main ICD10 (O682 – O68.2 Labour and delivery complicated by foetal heart rate anomaly with meconium in amniotic fluid)’]
20. (r =0.14) [‘Operation code (1512 – cone biopsy)’]
21. (r =0.14) [‘Treatment/medication code (1140871542 – mefenamic acid)’]
22. (r =−0.14) [‘Diagnoses – main ICD10 (O60 – O60 Preterm delivery)’]
23. (r =0.14) [‘Non-cancer illness code, self-reported (1522 – grave’s disease)”]
24. (r =0.14) [‘Treatment/medication code (1140857620 – depo-provera 50mg/1ml injection)’]
25. (r =0.13) [‘Operation code (1542 – laser treatment cervix)’]
26. (r =0.13) [‘Diagnoses – main ICD10 (J440 – J44.0 Chronic obstructive pulmonary disease with acute lower respiratory infection)’]
27. (r =0.13) [‘Treatment/medication code (1140883548 – ipratropium)’]
28. (r =0.13) [‘Diagnoses – main ICD10 (S430 – S43.0 Dislocation of shoulder joint)’]
29. (r =0.13) [‘Treatment/medication code (1140921822 – mirena 20mcg/24hrs intrauterine system)’]
30. (r =−0.13) [‘Diagnoses – main ICD10 (M819 – M81.9 Osteoporosis, unspecified)’]
31. (r =0.13) [‘Diagnoses – secondary ICD10 (M0690 – M06.90 Rheumatoid arthritis, unspecified (Multiple sites))’]
32. (r =0.13) [‘Diagnoses – main ICD10 (J441 – J44.1 Chronic obstructive pulmonary disease with acute exacerbation, unspecified)’]
33. (r =0.13) [‘Diagnoses – secondary ICD10 (N803 – N80.3 Endometriosis of pelvic peritoneum)’]
34. (r =0.13) [‘Treatment/medication code (1140871732 – buprenorphine)’]
35. (r =−0.13) [‘Histology of cancer tumour (8520 – Lobular carcinoma, NOS)’]
36. (r =−0.13) [‘Diagnoses – main ICD10 (O631 – O63.1 Prolonged second stage (of labour))’]
37. (r =−0.13) [‘Diagnoses – main ICD10 (O049 – O04.9 Complete or unspecified, without complication)’]
38. (r =−0.12) [‘Non-cancer illness code, self-reported (1606 – incisional hernia)’]
39. (r =0.12) [‘Diagnoses – secondary ICD10 (M205 – M20.5 Other deformities of toe(s) (acquired))’]
40. (r =0.12) [‘Treatment/medication code (1140867726 – lofepramine)’]
41. (r =0.12) [‘Diagnoses – secondary ICD10 (O16 – O16 Unspecified maternal hypertension)’]
42. (r =0.12) [‘Diagnoses – main ICD10 (N803 – N80.3 Endometriosis of pelvic peritoneum)’]
43. (r =0.12) [‘Treatment/medication code (1140884600 – metformin)’]
44. (r =0.12) [‘Diagnoses – main ICD10 (E042 – E04.2 Non-toxic multinodular goitre)’]
45. (r =0.12) [‘Diagnoses – secondary ICD10 (J069 – J06.9 Acute upper respiratory infection, unspecified)’]
46. (r =0.11) [‘Non-cancer illness code, self-reported (1556 – menorrhagia (unknown cause))’]
47. (r =0.11) [‘Diabetes diagnosed by doctor (2.0)’]
48. (r =−0.11) [‘Diagnoses – main ICD10 (Z400 – Z40.0 Prophylactic surgery for risk-factors related to malignant neoplasms)’]
49. (r =0.11) [‘Non-cancer illness code, self-reported (1220 – diabetes)’]
50. (r =0.11) [‘Non-cancer illness code, self-reported (1065 – hypertension)’]

